# Pleiotrophin deletion prevents high-fat diet-induced cognitive impairment, glial responses, and alterations of the perineuronal nets in the hippocampus

**DOI:** 10.1101/2024.09.23.614574

**Authors:** Héctor Cañeque-Rufo, Teresa Fontán-Baselga, Milagros Galán-Llario, Agata Zuccaro, María Gracia Sánchez-Alonso, Esther Gramage, María del Pilar Ramos-Álvarez, Gonzalo Herradón

**Affiliations:** Department of Health and Pharmaceutical Sciences, School of Pharmacy, Universidad San Pablo-CEU, CEU Universities, Urbanización Montepríncipe, 28660 Boadilla del Monte, Spain; Department of Chemistry and Biochemistry, School of Pharmacy, Universidad San Pablo-CEU, CEU Universities, Urbanización Montepríncipe, 28660 Boadilla del Monte, Spain

**Keywords:** Pleiotrophin, neuroinflammation, metabolic syndrome, perineuronal nets, obesity

## Abstract

Obesity and metabolic disorders, such as metabolic syndrome (MetS) facilitate the development of neurodegenerative diseases and cognitive decline. Persistent neuroinflammation plays an important role in this process. Pleiotrophin (PTN) is a cytokine that regulates energy metabolism and high-fat diet (HFD)-induced neuroinflammation, suggesting that PTN could play an important role in the connection between obesity and brain alterations, including cognitive decline. To test this hypothesis, we used an HFD-induced obesity model in *Ptn* genetically deficient mice (*Ptn*^−/−^). First, we confirmed that *Ptn* deletion prevents HFD-induced obesity. Our findings demonstrate that feeding wild-type (*Ptn*^+/+^) mice with HFD for 6 months results in short- and long-term memory loss in the novel object recognition task. Surprisingly, we did not observe any sign of cognitive impairment in *Ptn*^−/−^ mice fed with HFD. In addition, we observed that HFD induced microglial responses, astrocyte depletion, and perineuronal nets (PNNs) alterations in *Ptn*^+/+^ mice, while these effects of HFD were mostly prevented in *Ptn*^−/−^ mice. These results show a crucial role of PTN in metabolic responses and brain alterations induced by HFD and suggest the PTN signalling pathway as a promising therapeutic target for brain disorders associated with MetS.

## INTRODUCTION

During the last two decades, a significant increase in the prevalence of neurodegenerative diseases, such as Alzheimer’s disease (AD), has been observed. Despite extensive research, the specific factors that contribute to the development of these pathologies remain to be fully understood. Among them, obesity and related metabolic alterations, such as Metabolic Syndrome (MetS), a complex disorder characterised by cardiovascular and metabolic dysfunctions derived from obesity (Cornier et al., 2008), facilitate cognitive decline and are risk factors for neurodegenerative diseases, including AD (Castillo et al., 2019; Pugazhenthi et al., 2017; Rebelos et al., 2021; Santos et al., 2017).

The main connection between obesity, MetS, and neurodegenerative disorders is the persistent inflammation induced by metabolic alterations (de Bem et al., 2020). The consumption of a high-fat diet (HFD) leads to different changes linked to MetS, culminating in a systemic state of low-grade inflammation characterised by increased activity of peripheral pro-inflammatory cytokines in response to energy excess, known as metainflammation (Ali et al., 2020; Cavaliere et al., 2019; Dabke et al., 2019; Herradon et al., 2019). This is a factor that not only produces alterations at the peripheral organ level but is capable of producing numerous changes in the central nervous system (CNS), contributing to neuroinflammation and neuronal disruption, common features in multiple neurodegenerative diseases (Herradon et al., 2019).

This connection is supported by animal models with metabolic disorders showing numerous brain alterations associated with neurodegenerative diseases (Więckowska-Gacek et al., 2021). These models have identified persistent inflammation in metabolically active tissues as a trigger for neuroinflammation and neurodegeneration, and have unravelled some of the molecular mechanisms that regulate this connection, including the pleiotrophin (PTN) signalling pathway (Herradon et al., 2019).

PTN is a cytokine with a critical role in promoting the repair, survival, and differentiation of neurons in the CNS (Herradón & Pérez-García, 2014). PTN also acts as a potent regulator of neuroinflammation in different contexts (Cañeque-Rufo et al., 2023; Fernández-Calle et al., 2020; Fernández-Calle et al., 2017; Herradon et al., 2019; Vicente-Rodríguez et al., 2016). PTN interacts with different receptors in different organs, being Receptor Protein Tyrosine Phosphatase β/ζ (RPTPβ/ζ, also known as PTPRZ1) the most relevant in modulating the effects of PTN in CNS and neuroinflammation (Herradon et al., 2019). PTN binds RPTPβ/ζ (Herradón & Ezquerra, 2009), which is primarily expressed in the adult CNS in both neurons and glia (Herradón & Pérez-García, 2014; Maeda et al., 1994), and inhibits its phosphatase activity. This mechanism regulates the tyrosine phosphorylation of RPTPβ/ζ substrates involved in neuroinflammation (Herradon et al., 2019).

In previous studies, we have shown that PTN plays a key role in regulating insulin sensitivity, energy metabolism, and thermogenesis (Sevillano et al., 2019; Zuccaro et al., 2021). In addition, we have recently shown that endogenous PTN is necessary for the development of neuroinflammation, and mitochondrial alterations induced by HFD (Cañeque-Rufo et al., 2023). Altogether, these previous findings lead us to hypothesise that PTN plays an important role in the connection between MetS, obesity, cognitive impairment, and neurodegenerative diseases. To test this hypothesis, we used an HFD-induced obesity model in *Ptn* genetically deficient (*Ptn*^−/−^) mice. This study provides appreciable evidence that *Ptn* deletion protects against HFD-induced memory loss and glial responses. Furthermore, this study unravels for the first time that PTN is key in the correct formation and maintenance of perineuronal nets (PNNs), and the modulation of the alterations induced by HFD on PNNs. This is important because PNNs are a specialised extracellular matrix surrounding specific neurons (Härtig et al., 1999; Reichelt et al., 2019; Rossier et al., 2015). They are worthy of note for their synapse-stabilising role and for acting as a shield that protects neurons and synapses from potentially harmful neurochemical stimuli, such as oxidative stress and neuroinflammatory processes (Morawski et al., 2004; Reichelt et al., 2019; Suttkus et al., 2016), prominent features of the brain in a context of obesity and MetS (Cañeque-Rufo et al., 2023).

## MATERIALS AND METHODS

### Animals

*Ptn*^−/−^ mice were generated as previously described (Amet et al., 2001). C57BL/6J wild-type (*Ptn*^+/+^) and *Ptn*^−/−^ mice were divided randomly and housed in a specific pathogen-free room at 22 ± 1°C with 12h light/dark cycles, with free access to water and randomly fed with chow (Standard diet (STD), 18 kcal% fat, 58 kcal% carbohydrates, and 24% kcal protein; 3.1 kcal·g^−1^) or a High-Fat Diet (HFD, D12492, 60 kcal% fat, 20 kcal% carbohydrates, and 20% kcal protein; 5.24 kcal·g^−1^) as corresponding. Diet administration was started at 3 months old and maintained for 6 months (**Supplementary figure 1A**). Animals were fed *ad libitum*. All the animals were handled and maintained in accordance with the European Union Laboratory Animal Care Rules (2010/63/EU directive) and protocols were approved by the Animal Research Committee of CEU San Pablo University and by Comunidad de Madrid (PROEX 140.3/22).

### Novel Object Recognition test

At 22 weeks of treatment with the different diets, all mice were subjected to the novel object recognition (NOR) test (**Supplementary figure 1A**). The experimental apparatus consisted of a black quadrangular box (25 x 25 x 25 cm). The protocol of the experiment is summarized in **supplementary figure 1B**. Briefly, this test consists of four tasks, habituation, training, retention after 24 hours and retention after 5 days. In all the sessions, mice were placed in an equidistant position of both figures and were allowed to freely explore the apparatus and objects. For the habituation task, mice were allowed to explore freely for 10 min in the absence of objects. During the training phase, mice were placed in the experimental apparatus with two identical objects and allowed to explore for 10 min. After a retention interval of 24h, and 5 days, mice were placed again in the apparatus, in which one of the objects was replaced by a novel one in order to study the short- (retention interval of 24h) and long-term memory (retention interval of 5 days) (Antunes & Biala, 2012; Migues et al., 2016; Rodríguez-Matellán et al., 2020). Mice were allowed to explore for 10 min. Object exploration was defined as the orientation of the nose to the object at a distance ≤ 2 cm. All objects allow the mouse to stand on them, and they have the same height, width, and texture, ensuring that the animal explores the figures when they are currently novel. The object’s positions in the box (left or right) were chosen randomly. Preference for the novel object was expressed as the novel object exploration time percentage.

### Immunohistochemical analysis

#### Sacrifice and tissue sectioning

After 6h of fasting, 9-month-old mice from each experimental group (n = 6/group) were sacrificed by decapitation under CO_2_ exposure in gradual fill. The brains were removed from the skull and post-fixed overnight (O/N) in paraformaldehyde (PFA) 4% for 48h at 4°C and then kept in 30% sucrose. Coronal sections of 30 μm thickness were obtained in microtome (Leica SM2010 R).

#### Glial cells immunohistochemistry and image analysis

Randomly chosen hippocampal sections of the same coordinates range (from the Bregma -1.70 mm to -2.18 mm) were selected and incubated for 2 hours at RT in blocking solution (MgCl2 20 mM; 5% BSA; 10% NGS). Then, sections were incubated O/N at 4 °C with the following primary antibodies: mouse anti-glial fibrillary acidic protein (GFAP; Merck, Madrid, Spain; #MAB360; 1:1000) and rabbit anti-ionized calcium-binding adaptor molecule 1 (Iba1, Wako, Osaka, Japan; #019-19741; 1:1000). Next, the following secondary antibodies were incubated at a 1:500 concentration for 30 mins at RT to detect the binding of primary antibodies: Alexa-Fluor-488 donkey anti-rabbit (Invitrogen, Waltham, MA, USA; #A21206) and Alexa-Fluor-555 goat anti-mouse (Invitrogen, Waltham, MA, USA; #A21422). Finally, sections were mounted on positively charged glass slides. A commercial antifade mounting medium (Fluoromont^®^) was used to embed the sections.

Analysis was performed using *Fiji* software (NIH, Bethesda, MD, USA, Version 1.50f). Three images (10X objective) from every subject were acquired to estimate the number and size of Iba1^+^ and GFAP^+^ cells under an epifluorescence microscope (DM5500B, Leica, Solms, Germany) using the LAS X Core software (Leica Microsystems, Wetzlar, Germany; offline version). These images were processed using a previously described *Fiji* macro (Galán-Llario et al., 2023) aimed at enhancing cellular morphology detection and minimizing background artefacts. Next, the different parameters were tested using the “Analyze Particle” function previously described (Fernández-Calle et al., 2017). Additionally, to study the morphology of microglia more precisely, at least 30 Iba1^+^ cells in two images from different sections (40X objective) from each experimental condition were analyzed to determine the total process length and branching (Sholĺs analysis). All cells were traced using the *NeuronJ* plugin for *Fiji* and the total Iba1^+^ cells process length was obtained. Sholĺs analysis was used to analyse the complexity of the microglia processes and was performed using the plugin *ShollAnalysis* for *Fiji*. The Sholl’s Analysis plugin makes concentric circles, with a spacing of 1 μm between them, and marks the number of branches crossing each of these circles. After plotting the number of crossings against the distance to the microglia body, a representation of the complexity of the microglia processes was obtained.

### Perineuronal nets immunohistochemistry and image analysis

For the study of PNNs, we follow the protocol previously described (Galán-Llario et al., 2024). Briefly, randomly chosen hippocampal sections of the same coordinates range (from the Bregma -1.70 mm to -2.18 mm) were selected and blocked in a carbo-free blocking solution for 1 h at RT. Subsequently, slides were incubated O/N at 4 °C with biotinylated Wisteria Floribunda Agglutinin (WFA, Vector Laboratories, CA, USA; #B-1355-2 1:2000). Afterward, sections were incubated with Dylight 488-conjugated streptavidin antibody (Vector Laboratories, CA, USA; #SA-5488-1 1:400) for 1 h at RT. In the study of Parvalbumin (PV) positive neurons slides were incubated O/N at 4 °C with the primary antibody mouse anti-PV (Synaptic Systems, Göttingen, Germany; #195011; 1:1000) after WFA staining. Finally, slides were incubated at a 1:400 concentration for 1h at RT with the secondary antibody Alexa Fluor 555 donkey anti-mouse (Invitrogen, Waltham, MA, USA; #A21206). Finally, sections were mounted on positively charged glass slides. A commercial antifade mounting medium (Fluoromont^®^) was used to embed the sections.

Analysis was performed using two different software, *Fiji* (NIH, Bethesda, MD, USA, Version 1.50f) and Polygon AI (Polygon AI v3.0.3, Rewire Neuro, Inc.). Two images (10X objective) from every subject within the corresponding hippocampal area were acquired under an epifluorescence microscope (DM5500B, Leica, Solms, Germany) using the LAS X Core software (Leica Microsystems, Wetzlar, Germany; offline version). The conditions of a previously described analysis protocol were followed for the study of WFA fluorescence intensity of PNNs and to categorize PNNs into low, medium, and high intensity (Galán-Llario et al., 2024). For the categorisation of PNNs, quartiles were used as cutoff values based on the intensity of PNNs in images of *Ptn*^+/+^-STD mice within the corresponding area (<25% quartile for low-intensity; between 25% and 75% quartile for medium-intensity; >75% quartile for high-intensity). To estimate cell densities (WFA^+^ and PV^+^ cells/mm^3^), the area of the pyramidal cell layer (PCL), for CA areas, or granular cell layer (GCL), for dentate gyrus (DG), was measured in each image. These values were multiplied by the thickness of the slide to obtain the PCL or GCL volume (reference volume) of the image. Finally, the total number of cells positive for each marker was counted and divided by the reference volume. The values shown in the graphs correspond to the analysis of each image.

### Statistical analysis

Statistical analyses were performed using IBM-SPSS v25 software. The mean value and the standard error of the mean (±SEM) are represented in the graphs. The Shapiro-Wilk test was used for the normalities of the sample distribution. For the NOR test, body weight over time, food intake, and Sholl’s analysis data were analysed using a three-way repeated measures ANOVA test having sex, genotype, and diet as variables. To compare low-, medium-, and high-intensity PNNs, a two-tailed Fisher’s Exact test was performed to evaluate the distribution among experimental groups. For the rest of the studies, data were analysed using a three-way ANOVA test with the same variables. When relevant, to better dissect the effect of each variable, we used a two-way ANOVA, excluding the non-significant variable if the three-way ANOVA results allowed it. Bonferroni’s *Post-hoc* analysis was used to compare the differences between individual groups when the interaction between variables was found. All references to statistical significance are made to the three-way or two-way ANOVA test’s individual factors and their interaction. In the figures, significant differences revealed by three- and two-way ANOVAs and *Post-hoc* analyses are represented with (*) for differences between diets, (#) for differences between genotypes, and (&) for differences between sexes. A 95% confidence interval was used for statistical comparisons.

## RESULTS

### *Ptn* deletion prevents body weight gain induced by high-fat diet

First, we studied the weight gain induced by HFD in mice from the different experimental groups (**Figure 1A**). Three-way ANOVA revealed that both genotype and diet, but not sex, significantly affected the total increment of body weight (F_1, 110_ = 307,447; *p* < 0,001 (genotype); F_1, 110_ = 394,070; *p* < 0,001 (diet)). However, a significant interaction between the three variables was found (F_1, 110_ = 9,359; *p* = 0,003) (**Figure 1B**). We observed a very significant and nearly identical increase in body weight of male and female *Ptn*^+/+^ mice fed with HFD (**Figure 1B**). In contrast, we detected a very significant decrease in HFD-induced increment of body weight in *Ptn*^−/−^ mice compared with *Ptn*^+/+^ mice. In fact, we did not observe an HFD-induced significant increase in body weight in female *Ptn*^−/−^ mice (**Figure 1B**). Then, we studied the body weight gain during the weeks on diet. Two-way repeated measure ANOVA revealed that both genotype and diet significantly affected the body weight gain over time in both males (F_1, 51_ = 62,676; *p* < 0,001 (genotype); F_1, 51_ = 112,670; *p* < 0,001 (diet)) and females (F_1, 52_ = 73,482; *p* < 0,001 (genotype); F_1, 52_ = 107,444; *p* < 0,001 (diet)) (**Figures 1C and D**). In addition, a significant interaction between variables was found in both sexes (F_1, 51_ = 27,636; *p* < 0,001 (male); F_1, 52_ = 49,548; *p* < 0,001 (female)). Globally, the results confirmed that the body weight gain induced by HFD was drastically smaller in *Ptn*^−/−^ than in *Ptn*^+/+^ animals (**Figure 1B-D**). Finally, the study of food intake throughout the experiment showed no significant differences between the experimental groups (**Supplementary figure 2**).

**Figure 1.**
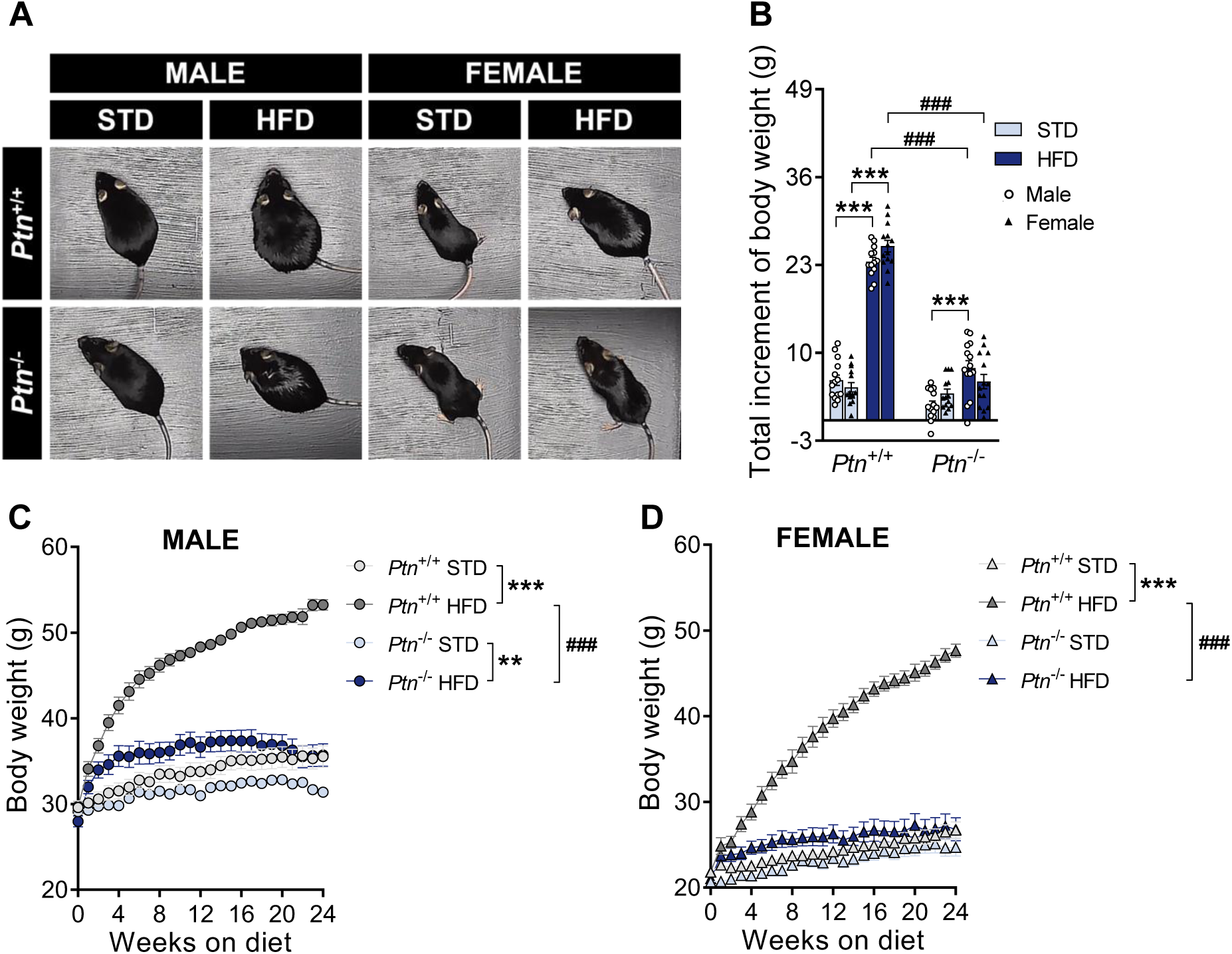
High-fat diet feeding and *Ptn* deletion effects on body weight. Representative images of the animals used (A); Total increment of body weight (B), and body weight over time in male (C) and female (D) *Ptn*^+/+^ and *Ptn*^−/−^ mice fed with a standard chow diet (STD) or with a high-fat diet (HFD) for 6 months. Data are presented as mean ± SEM (n = 14 animals per group; circles correspond to males and triangles correspond to females). \*\**p* < 0.01, \*\*\**p* < 0.001; ^###^*p* < 0.001.

### *Ptn* deletion protects against high-fat diet-induced memory loss

Using the NOR test, we analysed the short-term (24 h retention interval) and long-term (5 days retention interval) memory in animals from both genotypes fed with different diets. Three-way repeated measures ANOVA revealed that both genotype and diet, but not sex, significantly affected novel object recognition memory (F_1, 92_ = 197,714; *p* < 0,001 (genotype); F_1, 92_ = 124,983; *p* < 0,001 (diet)). To further explore the effect of genotype and diet, we employed two-way repeated measures ANOVA excluding the sex variable. Again, we found a significant effect of the genotype (F_1, 96_ = 185,541; *p* < 0,001) and diet (F_1, 96_ = 122,108; *p* < 0,001). In addition, a significant interaction between variables was found (F_1, 96_ = 124,053; *p* < 0,001). Our data revealed that *Ptn*^+/+^ animals fed with HFD show both short-term and long-term memory loss 24h and 5 days after the training session. However, *Ptn*^−/−^ mice fed with HFD did not show signs of memory loss in any retention task of the NOR test (**Figure 2**).

**Figure 2.**
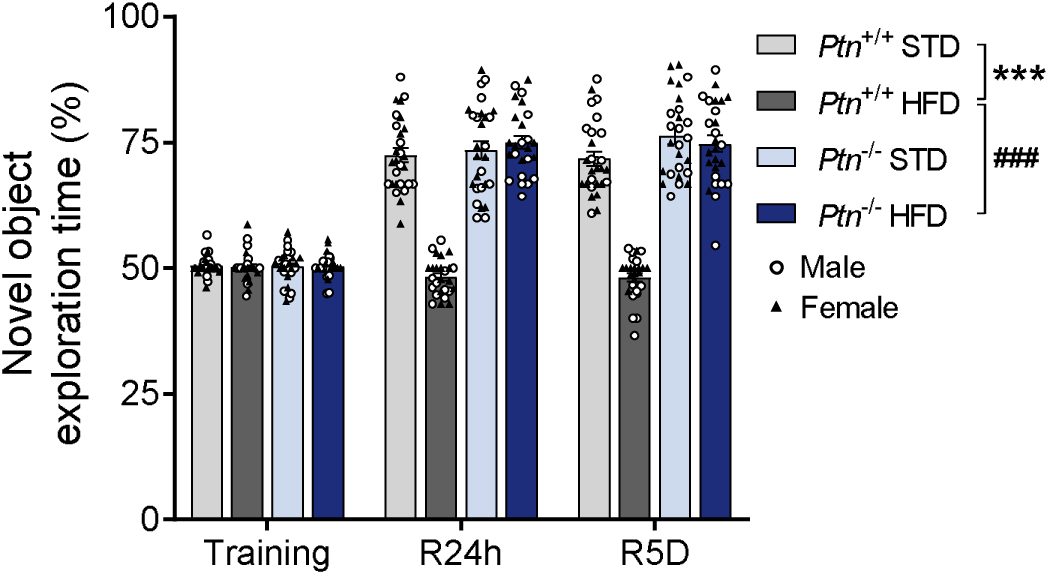
High-fat diet feeding and *Ptn* deletion effects on novel objects recognition memory. Percentage of time spent exploring the novel object in the different Novel Object Recognition (NOR) tasks (training, retention interval of 24 hours (R24h) and retention interval of 5 days (R5D)) by *Ptn*^+/+^ and *Ptn*^−/−^ mice fed with a standard chow diet (STD) or with a high-fat diet (HFD) for 6 months. Data are presented as mean ± SEM (n = 14 animals per group; circles correspond to males and triangles correspond to females). \*\*\**p* < 0.001; ^###^*p* < 0.001.

### *Ptn* deletion protects against astrocyte depletion and microglial changes induced by high-fat diet

We analysed the effects of HFD on glial cells in the hippocampus (**Figures 3 and 4**). Three-way ANOVA revealed that genotype and diet, but not sex, significantly affected the GFAP^+^ cells population (F_1, 47_ = 39,395; *p* < 0,001 (genotype); F_1, 47_ = 50,876; *p* < 0,001 (diet)) (**Figure 3B**). To further explore the effect of the genotype and diet, we employed two-way ANOVA excluding the sex variable, obtaining a significant effect of the genotype (F_1, 47_ = 42,310; *p* < 0,001), a significant effect of diet (F_1, 47_ = 54,641; *p* < 0,001), and a significant interaction between both variables (F_1, 47_ = 15,972; *p* < 0,001) (**Figure 3B**). The data showed that HFD decreased the hippocampal GFAP^+^ cell population in *Ptn*^+/+^ mice. Interestingly, this effect of HFD was not observed in *Ptn*^−/−^ mice (**Figure 3B**). The effect of HFD on the number of glial cells in this model seems to be specific to astrocytes, since we did not detect alterations in the number of microglial cells due to genotype or diet (**Figure 4B**). Following this observation, we analysed the morphology of microglial cells. Three-way ANOVA revealed that both genotype and diet, but not sex, significantly affected the average size of the Iba1^+^ cells (F_1, 47_ = 10,441; *p* = 0,002 (genotype); F_1, 47_ = 11,153; *p* = 0,002 (diet)) (**Figure 4C**). To further explore the effect of the genotype and diet, we employed two-way ANOVA excluding the sex variable, revealing a significant effect of the genotype (F_1, 47_ = 42,310; *p* < 0,001), a significant effect of diet (F_1, 47_ = 54,641; *p* < 0,001), and a significant interaction between both variables (F_1, 47_ = 15,972; *p* < 0,001). *Ptn*^+/+^ mice fed with HFD showed a significant increase in the average size of Iba1^+^ cells in the hippocampus. In contrast, this effect of HFD was not observed in *Ptn*^−/−^ mice (**Figure 4C**). Continuing with the morphometric analysis, three-way ANOVA revealed that genotype, diet, and sex significantly affected the body size (F_1, 323_ = 53,936; *p* > 0,001 (genotype); F_1, 323_ = 38,490; *p* > 0,001 (diet); (F_1, 323_ = 6,931; *p* = 0,009 (sex)) (**Figure 4D**) and ramification length (F_1, 323_ = 167,498; *p* > 0,001 (genotype); (F_1, 323_ = 136,677; *p* > 0,001 (diet); (F_1, 323_ = 4,983; *p* = 0,026 (sex)) (**Figure 4E**) of Iba1^+^ cells, although we did not detect significant interactions between variables. In general, we observed that female mice had a larger body size and shorter ramification length of Iba1^+^ cells compared to males regardless of genotype and diet. In addition, *Ptn*^+/+^ mice fed with HFD showed a population of Iba1^+^ cells with increased body size and decreased ramification length compared to *Ptn*^+/+^ mice that were fed an STD. Surprisingly, we observed that *Ptn*^−/−^ animals did not exhibit these HFD-induced effects in body size and ramification length of hippocampal Iba1^+^ cells (**Figure 4D and E**). Finally, three-way repeated measures ANOVA revealed that both genotype and diet, but not sex, significantly affected the microglia complexity measure obtained from Sholl’s analysis (F_1, 317_= 135,617; *p* > 0,001 (genotype); (F_1, 317_ = 126,195; *p* > 0,001 (diet)) (**Figure 4F**). To further explore the effect of the genotype and diet, we employed two-way repeated measures ANOVA excluding the sex variable. Again, we detected a significant effect of genotype (F_1, 321_= 135,347; *p* > 0,001), a significant effect of diet (F_1, 321_ = 125,106; *p* > 0,001), and a significant interaction between both variables (F_1, 321_ = 71,108; *p* < 0,001) (**Figure 4F**). We found that HFD induced a decrease in the complexity of Iba1^+^ cells in the hippocampus of *Ptn*^+/+^ mice, an effect that was not observed in *Ptn*^−/−^ mice fed with HFD (**Figure 4F**).

**Figure 3.**
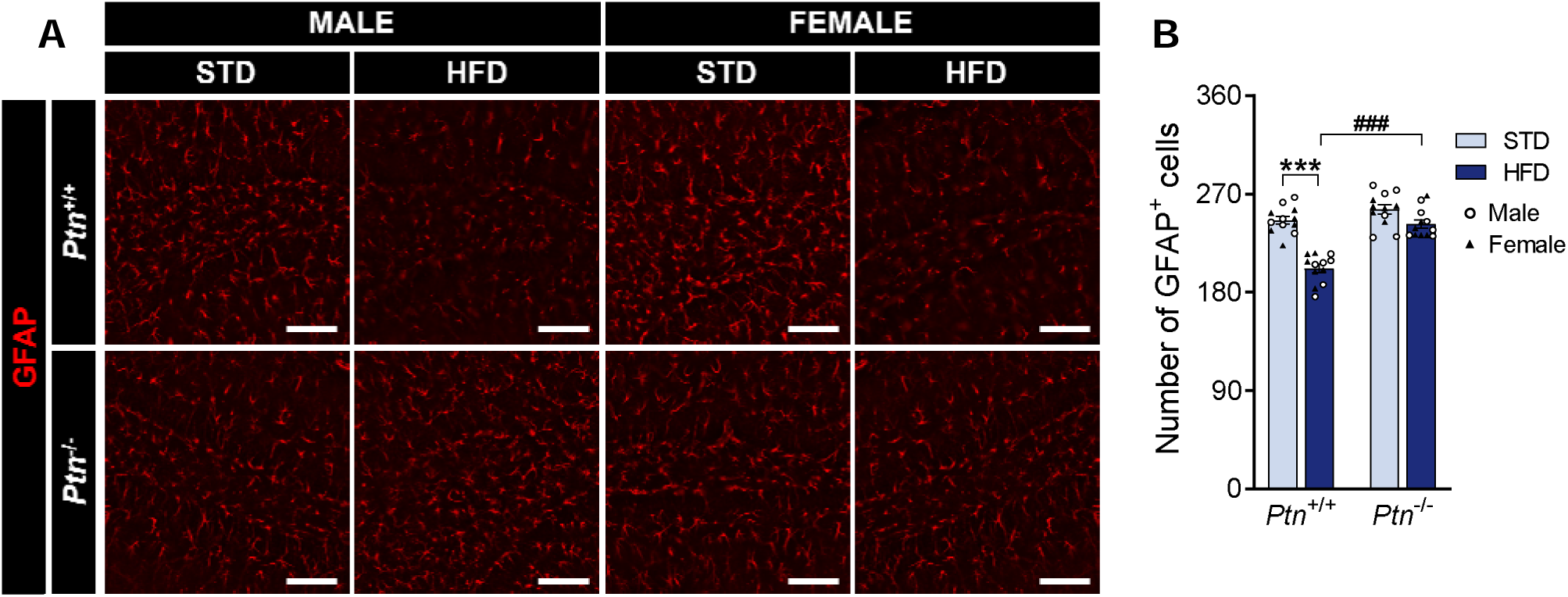
High-fat diet feeding and *Ptn* deletion effects on astrocytes. Representative images showing GFAP fluorescence from the different experimental groups (A). Quantification of the number of GFAP^+^ cells per field (B) in the hippocampus of *Ptn*^+/+^ and *Ptn*^−/−^ mice fed with a standard chow diet (STD) or with a high-fat diet (HFD). Data are presented as mean ± SEM (n = 6 animals per group; circles correspond to males and triangles correspond to females). \*\*\**p* < 0.001; ^###^*p* < 0.001. Scale bar 100 µm.

**Figure 4.**
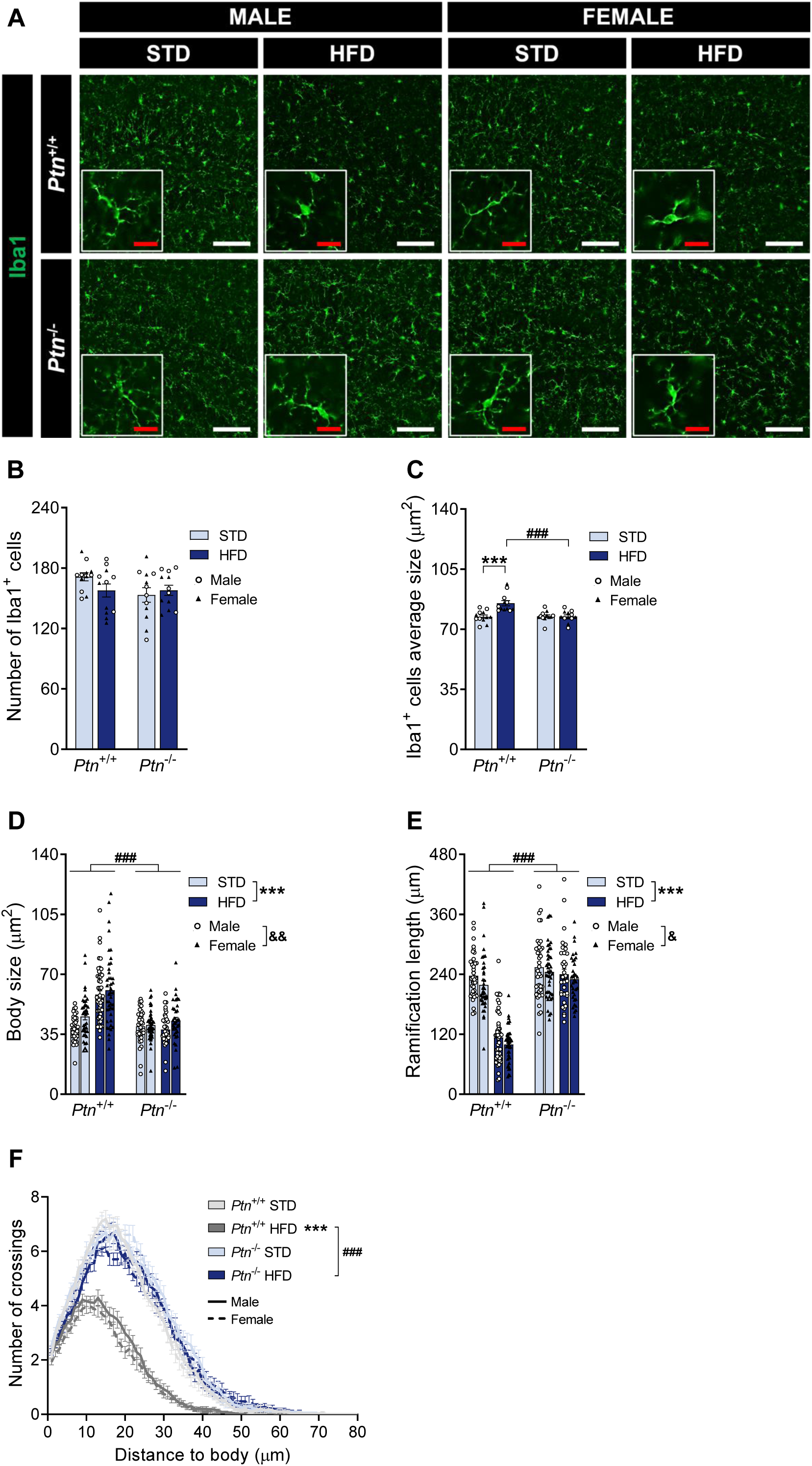
High-fat diet feeding and *Ptn* deletion effects on microglia polarisation. Representative images showing Iba1 fluorescence from the different experimental groups (A). Quantification of the number of Iba1^+^ cells per field (B); average size (C); body size (D), ramification length (E), and complexity (F) from Iba1^+^ cells in the hippocampi of male and female *Ptn*^+/+^ and *Ptn*^−/−^ mice fed with a standard chow diet (STD) or with a high-fat diet (HFD). Data are presented as mean ± SEM (n = 6 animals per group; circles correspond to males and triangles correspond to females). \*\*\**p* < 0.001; ^###^*p* < 0.001; ^&^*p* < 0.05, ^&&^*p* < 0.01. White scale bar 100 µm. Red scale bar 20 µm.

### High-fat diet and *Ptn* deletion alter the number and intensity of perineuronal nets and the number of PV^+^ cells in the dentate gyrus

We studied PNNs in the different hippocampal regions. We started analysing WFA^+^ and PV^+^ cells in the DG of mice belonging to the different experimental groups (**Figure 5A**). Three-way ANOVA revealed that genotype, diet, and sex, significantly affect the number of WFA^+^ cells in the DG (F_1, 95_ = 65,496; *p* < 0,001 (genotype); (F_1, 95_ = 7,079; *p* = 0,009 (diet); (F_1, 95_ = 8,047; *p* = 0,006 (sex)). However, no significant interaction between variables was found (**Figure 5B**). Overall, we observed that female mice have a lower number of WFA^+^ cells in the DG compared to males regardless of genotype and diet. In addition, HFD consumption increased the number of WFA^+^ cells in the DG of *Ptn*^+/+^ mice. Interestingly, we observed that *Ptn*^−/−^ mice have a lower number of WFA^+^ cells, which were not affected by HFD (**Figure 5B**). Three-way ANOVA revealed that both genotype and sex, but not diet, significantly affect the WFA^+^ cells intensity in the DG (F_1, 95_ = 16,375; *p* < 0,001 (genotype); (F_1, 95_ = 5,937; *p* = 0,017 (sex)), although no significant interaction was detected between variables (**Figure 5C**). Female mice showed higher WFA^+^ cell intensity in the DG compared to males regardless of genotype and diet. Moreover, we observed an increase in WFA^+^ cell intensity in *Ptn*^−/−^ mice regardless of the diet consumed (**Figure 5C**). This is related to the significant increase in high-intensity WFA^+^ cells percentage found in *Ptn*^−/−^ mice fed with HFD compared to *Ptn*^+/+^ mice and the general decrease in low-intensity WFA^+^ cells percentage of *Ptn*^−/−^ mice (**Figure 5D**).

**Figure 5.**
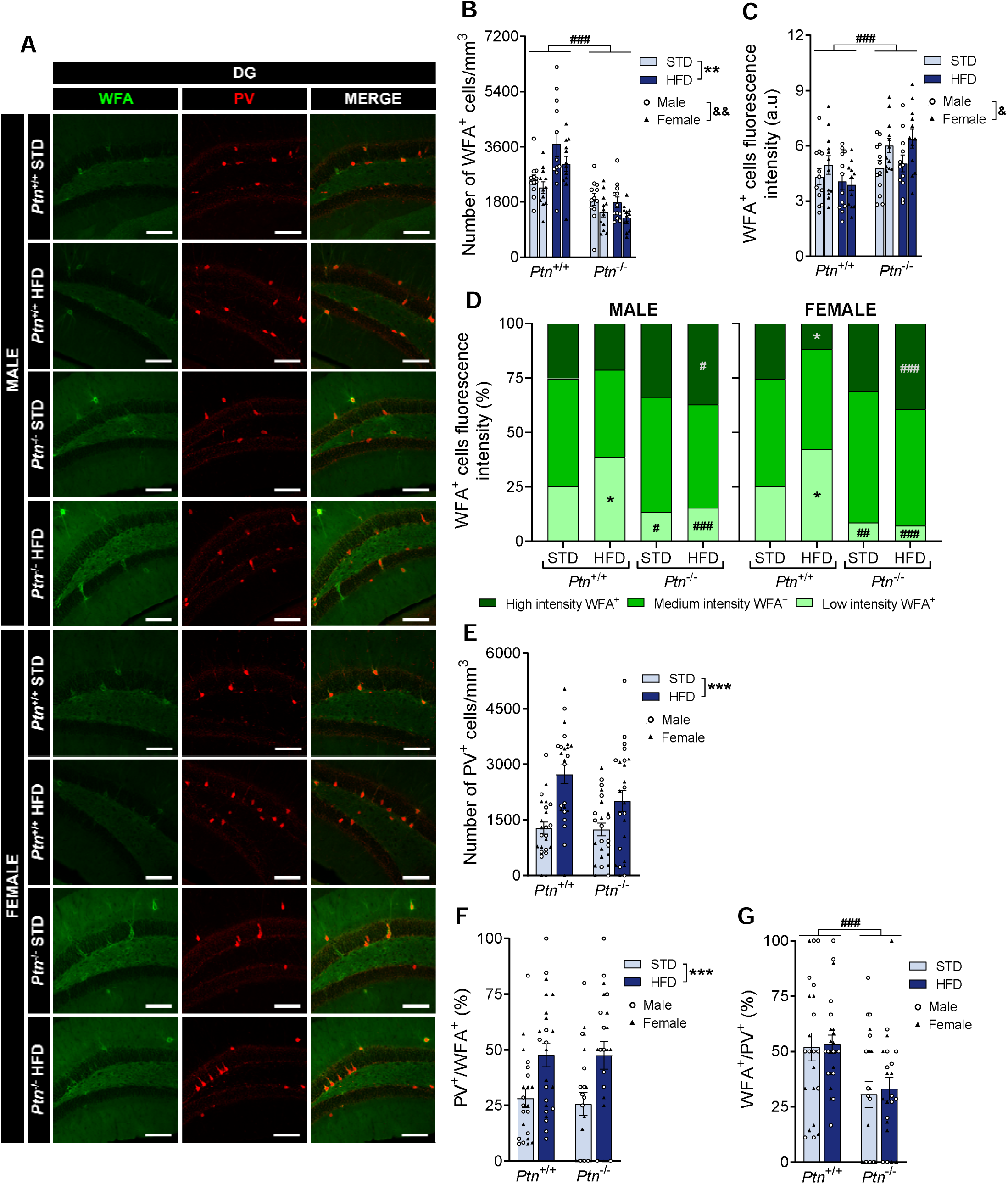
High-fat diet feeding and *Ptn* deletion effects on perineuronal nets and parvalbumin^+^ cells in the dentate gyrus. Representative images showing WFA and PV fluorescence from the different experimental groups (A). Quantification of the number of WFA^+^ cells per mm^3^ (B); WFA^+^ cells fluorescence intensity (C); percentage of WFA^+^ cells corresponding to different WFA intensity levels (D); number of PV^+^ cells per mm^3^ (E); percentage of WFA^+^ cells surrounding PV^+^ cells (F), and percentage of PV^+^ cells that are surrounded by WFA^+^ cells (G) in the DG of *Ptn*^+/+^ and *Ptn*^−/−^ mice fed with a standard chow diet (STD) or with a high-fat diet (HFD). Data are presented as mean ± SEM (n = 6 animals per group; circles correspond to males and triangles correspond to females). \**p* < 0.05, \*\**p* < 0.01, \*\*\**p* < 0.001; ^#^*p* < 0.05, ^##^*p* < 0.01, ^###^*p* < 0.001; ^&^*p* < 0.05, ^&&^*p* < 0.01. Scale bar 100 µm.

We also studied PV^+^ cells in the DG. Three-way ANOVA revealed that HFD significantly increased the number of these cells (F_1, 95_ = 28,479; *p* < 0,001) independently of the sex and genotype (**Figure 5E**). In the colocalization analysis, three-way ANOVA revealed that HFD significantly increased the percentage of PNNs surrounding PV^+^ cells (F_1, 89_ = 15,269; *p* < 0,001) in both *Ptn*^+/+^ and *Ptn*^−/−^ mice regardless of sex (**Figure 5F**). Moreover, three-way ANOVA revealed that deletion of *Ptn* significantly decreases the percentage of PV^+^ cells that are surrounded by PNNs (F_1, 89_ = 15,269; *p* < 0,001) regardless of diet and sex (**Figure 5G**).

### High-fat diet and *Ptn* deletion alter the number and intensity of perineuronal nets and the number of PV^+^ cells in the CA1 hippocampal region

We analysed WFA^+^ and PV^+^ cells in the CA1 hippocampal region (**Figure 6A**). Three-way ANOVA revealed that both genotype and diet, but not sex, significantly affect the number of WFA^+^ cells in the CA1 hippocampal region (F_1, 95_ = 80,511; *p* < 0,001 (genotype); (F_1, 95_ = 42,681; *p* < 0,001 (diet)) (**Figure 6B**). To further explore the effect of genotype and diet, we employed two-way ANOVA excluding the sex variable. Again, we detected a significant effect of genotype (F_1, 95_ = 80,878; *p* < 0,001), a significant effect of diet (F_1, 95_ = 42,875; *p* < 0,001) and a significant interaction between both variables (F_1, 95_ = 25,777; *p* < 0,001) (**Figure 6B**). We observed that HFD consumption decreased the number of WFA^+^ cells in the CA1 hippocampal region of *Ptn*^+/+^ mice. Interestingly, we observed that *Ptn*^−/−^ mice have a higher number of WFA^+^ cells, which were not affected by HFD (**Figure 6B**). Three-way ANOVA revealed a significant increase in WFA^+^ cells intensity of CA1 hippocampal region in *Ptn*^−/−^ mice regardless of sex and diet consumed (F_1, 95_ = 201,266; *p* < 0,001) (**Figure 6C**). This is related to the significant increase in high-intensity WFA^+^ cells percentage found in *Ptn*^−/−^ mice compared to the *Ptn*^+/+^ mice and to the general decrease in medium- and low-intensity WFA^+^ cells percentage of *Ptn*^−/−^ mice (**Figure 6D**).

**Figure 6.**
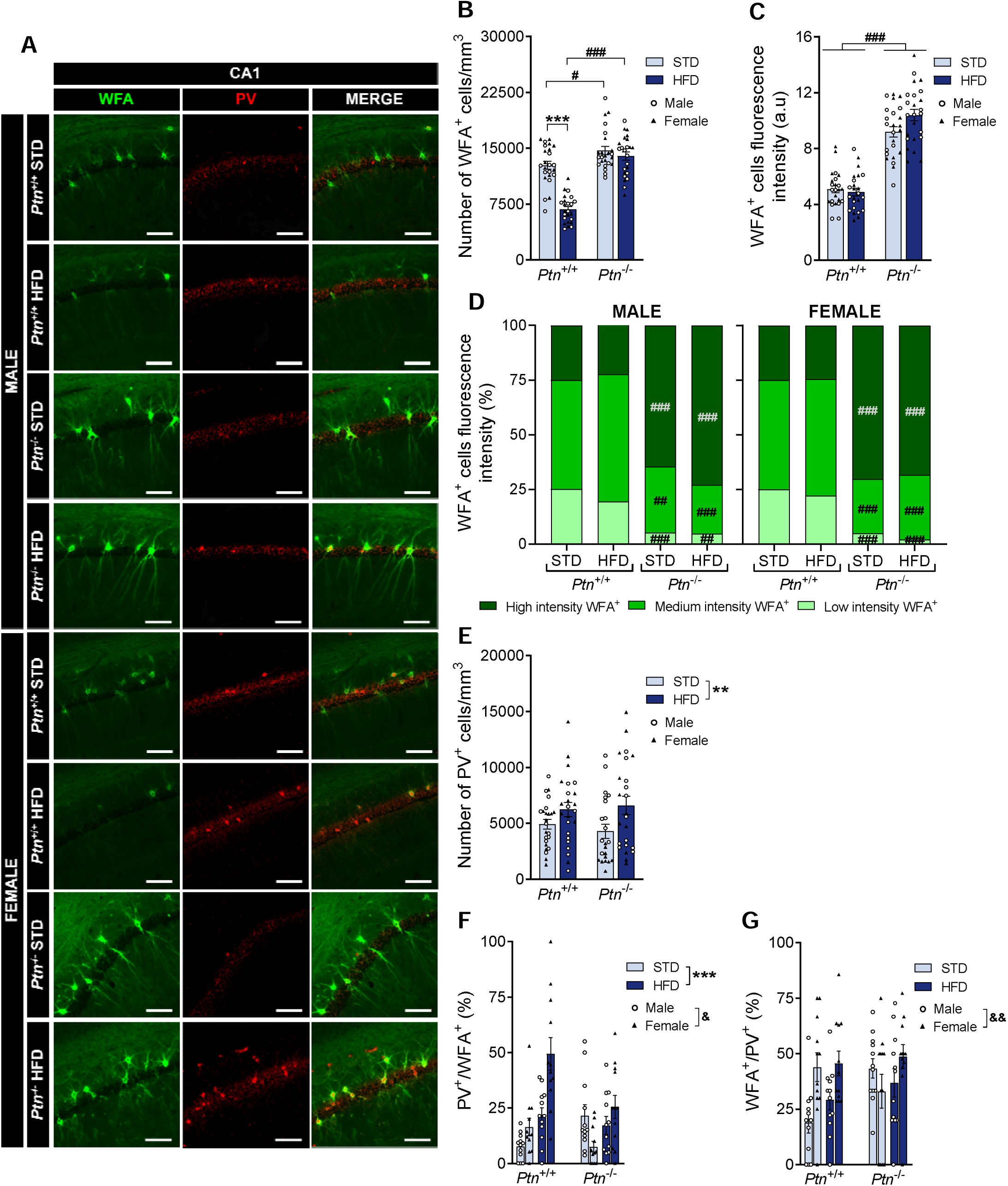
High-fat diet feeding and *Ptn* deletion effects on perineuronal nets and parvalbumin^+^ cells in the CA1 hippocampal region. Representative images showing WFA and PV fluorescence from the different experimental groups (A). Quantification of the number of WFA^+^ cells per mm^3^ (B); WFA^+^ cells fluorescence intensity (C); percentage of WFA^+^ cells corresponding to different WFA intensity levels (D); number of PV^+^ cells per mm^3^ (E); percentage of WFA^+^ cells surrounding PV^+^ cells (F), and percentage of PV^+^ cells that are surrounded by WFA^+^ cells (G) in the CA1 hippocampal region of *Ptn*^+/+^ and *Ptn*^−/−^ mice fed with a standard chow diet (STD) or with a high-fat diet (HFD). Data are presented as mean ± SEM (n = 6 animals per group; circles correspond to males and triangles correspond to females). \*\**p* < 0.01, \*\*\**p* < 0.001; ^#^*p* < 0.05, ^##^*p* < 0.01, ^###^*p* < 0.001; ^&^*p* < 0.05, ^&&^*p* < 0.01. Scale bar 100 µm.

We also studied PV^+^ cells in the CA1 hippocampal region. Three-way ANOVA revealed that HFD significantly increased the number of these cells (F_1, 93_ = 10,184; *p* = 0,002) in both *Ptn*^+/+^ and *Ptn*^−/−^ mice regardless of sex (**Figure 6E**). In the colocalization analysis, three-way ANOVA revealed that diet and sex, but not genotype, significantly affect the percentage of PNNs surrounding PV^+^ cells (F_1, 95_ = 23,331; *p* < 0,001 (diet); F_1, 95_ = 6,023; *p* = 0,016 (sex)), although no significant interaction was detected between variables (**Figure 6F**). Female *Ptn*^+/+^ mice showed a higher percentage of PNNs surrounding PV^+^ cells in the CA1 hippocampal region compared to males, especially in mice fed with a HFD (**Figure 6F**). In addition, three-way ANOVA revealed that female mice present a percentage of PV^+^ cells that are surrounded by PNNs significantly higher than male mice (F_1, 95_ = 7,090; *p* = 0,009) regardless of diet and genotype (**Figure 6G**).

### High-fat diet and *Ptn* deletion alter the intensity of perineuronal nets and the number of PV^+^ cells in the CA2 hippocampal region

Next, we analysed the CA2 hippocampal region. However, the diffuse structuring of PNNs in this area makes it difficult to study them in an individualised manner (**Figure 7A**). For this reason, we studied only the WFA^+^ fluorescence intensity in this region. Three-way ANOVA revealed a significant increase in WFA^+^ intensity of CA2 hippocampal region in *Ptn*^−/−^ mice regardless of sex and diet consumed (F_1, 94_ = 2505,812; *p* < 0,001)) (**Figure 7B**).

**Figure 7.**
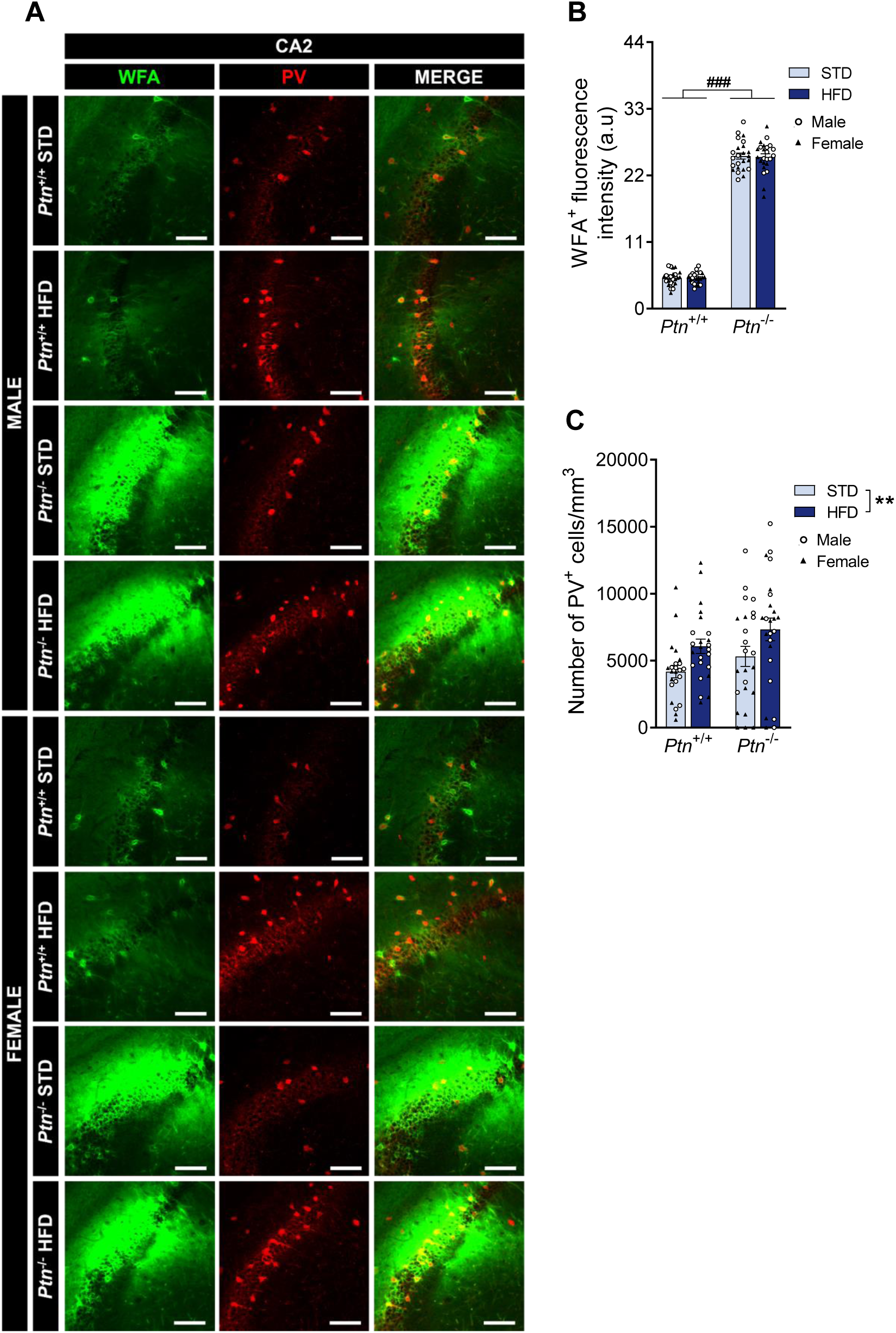
High-fat diet feeding and *Ptn* deletion effects on Perineuronal nets and Parvalbumin cells in the CA2 hippocampal region. Representative images showing WFA and PV fluorescence from the different experimental groups (A). Quantification of the WFA fluorescence intensity (B), and number of PV^+^ cells per mm^3^ (C) in the CA2 hippocampal region of *Ptn*^+/+^ and *Ptn*^−/−^ mice fed with a standard chow diet (STD) or with a high-fat diet (HFD). Data are presented as mean ± SEM (n = 6 animals per group; the white circles correspond to male data, while the black triangles correspond to female data). \*\**p* < 0.01; ^###^*p* < 0.001. Scale bar 100 µm.

We also studied PV^+^ cells in the CA2 hippocampal region. Three-way ANOVA revealed that HFD significantly increased the number of these cells (F_1, 94_ = 9,510; *p* = 0,003) independently of the sex and genotype (**Figure 7C**).

### High-fat diet and *Ptn* deletion alter the number and intensity of perineuronal nets and the number of PV^+^ cells in the CA3 hippocampal region

Finally, we analysed WFA^+^ and PV^+^ cells in the CA3 hippocampal region (**Figure 8A**). Three-way ANOVA revealed that both genotype and diet, but not sex, significantly affect the number of WFA^+^ cells in the CA3 hippocampal region (F_1, 95_ = 105,677; *p* < 0,001 (genotype); F_1, 95_ = 46,331; *p* < 0,001 (diet)) (**Figure 8B**). To further explore the effect of the genotype and diet, we employed two-way ANOVA excluding the sex variable. Again, we detected a significant effect of the genotype (F_1, 95_ = 103,933; *p* < 0,001), a significant effect of diet (F_1, 95_ = 45,566; *p* < 0,001) and a significant interaction between both variables (F_1, 95_ = 34,966; *p* < 0,001) (**Figure 8B**). Overall, we observed that HFD consumption decreased the number of WFA^+^ cells in the CA3 hippocampal region of *Ptn*^+/+^ mice. Interestingly, we observed that *Ptn*^−/−^ mice showed a higher number of WFA^+^ cells, which were not affected by HFD (**Figure 8B**). Three-way ANOVA revealed a significant increase in WFA^+^ cells intensity of CA3 hippocampal region in *Ptn*^−/−^ mice regardless of sex and diet consumed (F_1, 95_ = 186,351; *p* < 0,001) (**Figure 8C**). This is related to the significant increase in high-intensity WFA^+^ cell percentage found in *Ptn*^−/−^ mice compared to the *Ptn*^+/+^ mice. In addition, it is related to the significant decrease in medium-intensity WFA^+^ cells percentage found in male *Ptn*^−/−^ mice compared to *Ptn*^+/+^ mice and to the general decrease in low-intensity WFA^+^ cells percentage of *Ptn*^−/−^ mice (**Figure 8D**).

**Figure 8.**
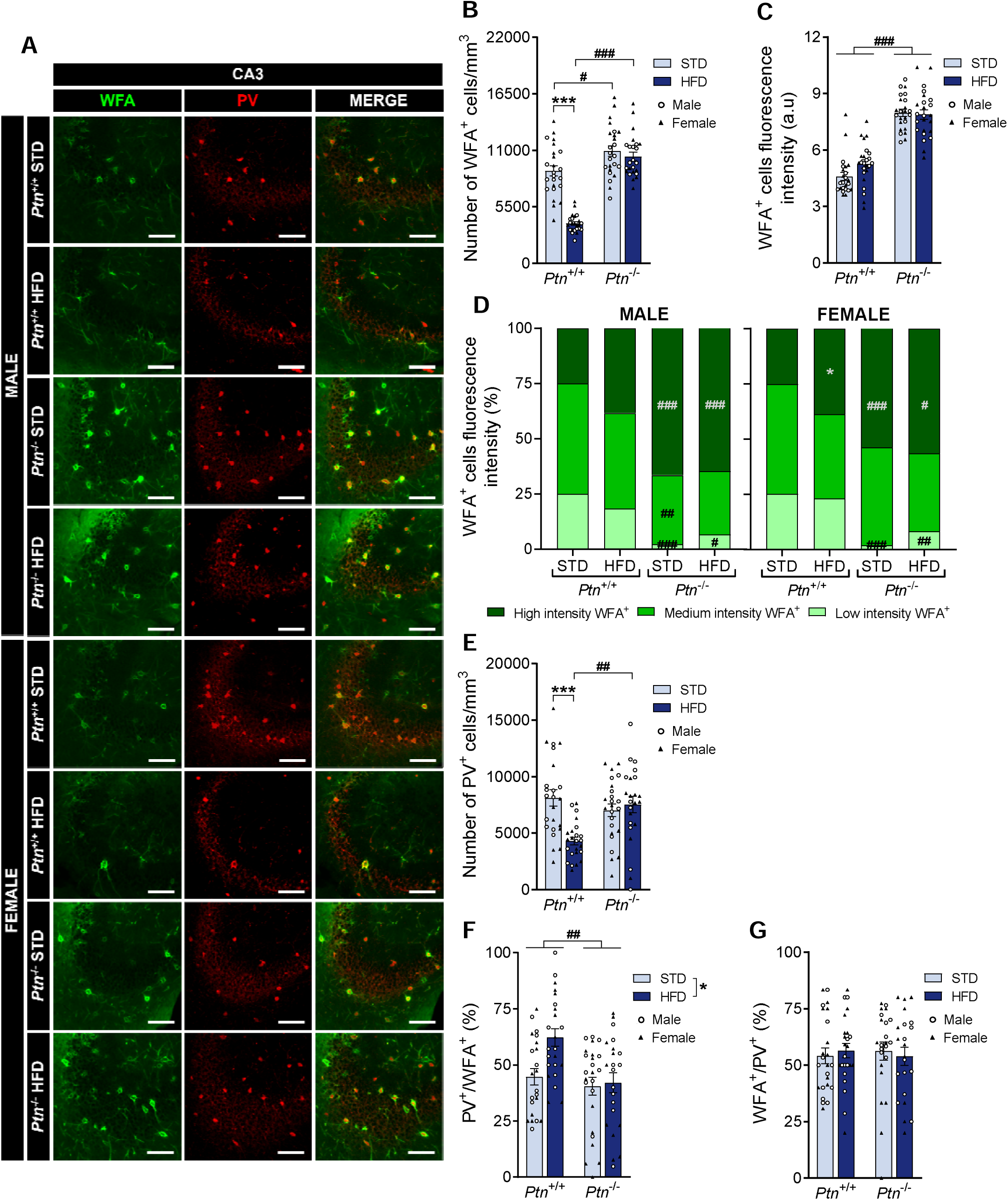
High-fat diet feeding and *Ptn* deletion effects on perineuronal nets and parvalbumin^+^ cells in the CA3 hippocampal region. Representative images showing WFA and PV fluorescence from the different experimental groups (A). Quantification of the number of WFA^+^ cells per mm^3^ (B); WFA^+^ cells fluorescence intensity (C); percentage of WFA^+^ cells corresponding to different WFA intensity levels (D); number of PV^+^ cells per mm^3^ (E); percentage of WFA^+^ cells surrounding PV^+^ cells (F), and percentage of PV^+^ cells that are surrounded by WFA^+^ cells (G) in the CA3 hippocampal region of *Ptn*^+/+^ and *Ptn*^−/−^ mice fed with a standard chow diet (STD) or with a high-fat diet (HFD). Data are presented as mean ± SEM (n = 6/group; circles correspond to males and triangles correspond to females). \**p* < 0.05, \*\*\**p* < 0.001; ^#^*p* < 0.05, ^##^*p* < 0.01, ^###^*p* < 0.001. Scale bar 100 µm.

We also studied PV^+^ cells in the CA3 hippocampal region. Three-way ANOVA revealed that only diet significantly affects the number of PV^+^ cells (F_1, 95_ = 8,184; *p* = 0,005). To further explore the effect of the genotype and diet, we employed two-way ANOVA excluding the sex variable. We detected a significant effect of diet (F_1, 95_ = 8,369; *p* = 0,005) and a significant interaction between genotype and diet (F_1, 95_ = 10,929; *p* = 0,001) (**Figure 8E**). We observed that HFD decreased the number of PV^+^ cells in the CA3 hippocampal region of *Ptn*^+/+^ mice. Interestingly, this HFD-induced decrease was not observed in *Ptn*^−/−^ mice (**Figure 8E**). In the colocalization analysis, three-way ANOVA revealed that genotype and diet, but not sex, significantly affect the percentage of PNNs surrounding PV^+^ cells (F_1, 91_ = 7,232; *p* = 0,009 (genotype); F_1, 91_ = 4,188; *p* = 0,044 (diet)) (**Figure 8F**). To further explore the effect of the genotype and diet, we employed two-way ANOVA excluding the sex variable. Again, we detected a significant effect of genotype (F_1, 91_ = 7,203; *p* = 0,009) and diet (F_1, 91_ = 4,171; *p* = 0,044), although no significant interaction was detected between variables (**Figure 8F**). HFD consumption increased the percentage of PNNs surrounding PV^+^ cells in the CA3 hippocampal region in *Ptn*^+/+^ mice. Moreover, *Ptn* deletion decreased the percentage of PNNs surrounding PV^+^ cells regardless of the diet and sex of the animals (**Figure 8F**). In addition, three-way ANOVA did not reveal significant differences in the percentage of PV^+^ cells that are surrounded by PNNs (**Figure 8G**).

## DISCUSSION

During the last few years, considerable progress has been made in the study of the association between MetS and the subsequent development of neurodegenerative pathologies like different dementias, including AD (de Bem et al., 2020; Ng et al., 2016). Early interventions targeting non-genetic risk factors are crucial for both prevention and treatment of these incapacitating disorders, as most cases correspond to the sporadic form of these diseases. The molecular mechanisms connecting MetS and neurodegeneration are not yet fully clear (Picone et al., 2020), but chronic inflammation in adipose and metabolic tissues seems to play a pivotal role (Więckowska-Gacek et al., 2021). Animal models of metabolic disorders share most of the brain alterations associated with neurodegenerative diseases, even in early life (Więckowska-Gacek et al., 2021). In the context of the events contributing to neurodegeneration, this provides an interesting tool to study the molecular mechanisms known to be involved in the modulation of both metabolic disorders and neuroinflammation, such as the PTN signalling pathway (Cañeque-Rufo et al., 2023; Fernández-Calle et al., 2017; Sevillano et al., 2019; Zuccaro et al., 2021).

As we observed in previous studies, our findings demonstrate that *Ptn* deletion prevents HFD-induced increase in body weight (Cañeque-Rufo et al., 2023; Zuccaro et al., 2021), revealing that this protective effect is consistent in both male and female mice. The data suggest a crucial role of PTN in regulating metabolic responses to HFD irrespective of sex. Moreover, the results demonstrate for the first time that *Ptn* deletion protects against the cognitive impairment induced by HFD. We observed that *Ptn*^+/+^ mice fed with HFD were unable to identify the novel object after a short (24 h) and a long retention interval (5 days). Strikingly, *Ptn*^−/−^ mice fed with HFD recalled the novel object after both retention intervals, suggesting a memory formation process unaffected by HFD. The data indicate that *Ptn* deletion fully prevents HFD-induced memory impairment.

Consumption of HFD also caused different alterations in the brain. We observed that HFD induced an astrocyte depletion in *Ptn*^+/+^ mice, which could lead to abnormalities in synaptic function and in the regulation of inflammatory responses. However, we observed that HFD-induced decrease in astrocytes was prevented in *Ptn*^−/−^ mice. Interestingly, morphometric analysis of microglia revealed that HFD-induced changes in these cells are associated with a polarisation of microglia towards a proinflammatory reactive phenotype in *Ptn*^+/+^ mice, which is consistent with our previous studies showing that HFD increases the expression of proinflammatory factors only in *Ptn*^+/+^ mice, not in *Ptn*^−/−^ mice (Cañeque-Rufo et al., 2023). Taken together, the data demonstrate that HFD leads to a PTN-dependent state of chronic neuroinflammation and glial responses.

The changes induced by HFD in glial cells suggest a profound effect of this diet in the brain environment, in which PNNs are important components. We found that HFD decreased PNNs in the CA regions of the hippocampus in *Ptn*^+/+^ mice. However, we observed an opposite response in the DG. PNNs play a crucial role in stabilizing synaptic connections, modulating neuronal excitability, and in neuronal protection and plasticity (Morawski et al., 2004; Reichelt et al., 2019; Suttkus et al., 2016). In the hippocampus, PNNs play a fundamental role in neurogenesis and learning (Cope & Gould, 2019; Crapser et al., 2020), suggesting that these HFD-induced changes in the PNNs of the hippocampus are related to the observed cognitive impairments observed in *Ptn*^+/+^ mice fed with this diet. Interestingly, *Ptn*^−/−^ mice fed with a standard diet showed a general increase of PNNs in the hippocampus compared with *Ptn*^+/+^ mice fed with STD. More importantly, prolonged HFD consumption did not cause learning deficits in *Ptn*^−/−^ mice, which correlated with the absence of significant alterations in hippocampal PNNs induced by HFD in these mice. Taken together, the data suggest that PTN plays a fundamental role in the formation and maintenance of PNNs and in the regulation of HFD-induced changes in PNNs and cognitive impairment.

Previous studies have identified the PTN receptor, RPTPβ/ζ, as a crucial element of PNNs by participating in their formation and stabilization (Eill et al., 2020; Sinha et al., 2023). On the other hand, matrix metalloproteinase 9 (MMP9) is the primary enzyme responsible for PNNs degradation (Gray et al., 2008; Reichelt et al., 2019), and its reduced activity promotes the formation of PNNs (Wen et al., 2018). HFD consumption leads to an increase in the activity of this enzyme, potentially resulting in increased degradation of PNNs under obesity conditions (Reichelt et al., 2019). Previously, we showed that *Ptn*^−/−^ mice have an increased cerebral expression of *Ptprz1* and display nearly undetectable levels of *Mmp9* expression in the CNS (Cañeque-Rufo et al., 2023). All in all, the data suggest enhanced anchoring and stabilization of PNNs in *Ptn*^−/−^ mice caused by higher levels of expression of *Ptprz1*, and reduced degradation of PNNs due to *Mmp9* downregulation. Accordingly, in the present work, we demonstrate that *Ptn*^−/−^ mice show a higher number and intensity of PNNs, independently of the diet consumed. In addition, there is extensive literature defining that PNNs are found surrounding the neuron soma and proximal dendrites (Cope & Gould, 2019). This particular structuring results from pruning processes conducted by microglial cells (Crapser et al., 2020; Liu et al., 2024). Surprisingly, our study shows that *Ptn*^−/−^ animals have PNNs that are not exclusively restricted to the soma and proximal dendrites but are also found surrounding distal dendrites. This suggests that *Ptn* deletion leading to a state of generalised decreased inflammatory processes (Cañeque-Rufo et al., 2023) and reduced microglial activity may prevent the microglial-induced pruning of the PNNs.

PNNs primarily envelop fast-spiking neurons, facilitating the swift ionic exchange required for their rapid response (Tewari et al., 2022). Among these, PV^+^ interneurons, crucial for regulating neuron activity in the hippocampus, are frequently surrounded by PNNs (Jakovljević et al., 2021). We observed that HFD produced an increase in the number of PV^+^ interneurons in the CA1, CA2 and DG regions in both *Ptn*^+/+^ and *Ptn*^−/−^ mice. However, HFD decreased the number of PV^+^ interneurons only in the CA3 area of *Ptn*^+/+^ mice, suggesting a region-specific modulation of PV^+^ interneurons in the mouse hippocampus, possibly related to the different connections and circuits involved in each region. In addition, we observed that the percentage of PNNs surrounding PV^+^ interneurons increased after HFD consumption. However, the percentage of PV^+^ interneurons that are surrounded by PNNs was unaltered, indicating that the loss of PNNs induced by HFD involves mostly PNNs that do not surround PV^+^ interneurons. Interestingly, in a single-cell study (Arneson et al., 2019), it was shown that HFD consumption produces an overexpression of *Ptprz1* (RPTPβ/ζ) only in GABAergic neurons (**Supplementary Figure 3**), which would potentially lead to enhanced anchoring of PNNs in GABAergic neurons, making them more resistant to the effects of HFD.

## CONCLUSIONS

The findings presented here provide substantial evidence that *Ptn* deletion protects against HFD-induced obesity. The data demonstrate that the absence of endogenous PTN fully prevents the cognitive impairment induced by prolonged consumption of HFD, which may be related to the attenuation of HFD-induced alterations of hippocampal PNNs, and glial responses caused by *Ptn* deletion.

The evidence presented here suggests that targeting PTN may be a novel therapeutic strategy to disrupt the connection between MetS, obesity, neuroinflammation and cognitive decline.

## Data availability

Multitissue Single-Cell Analysis Reveals Differential Tissue, Cellular, and Molecular Sensitivity Between Fructose and High Fat High Sucrose Diets (Arneson et al., 2019) published on Single Cell Portal (https://singlecell.broadinstitute.org/single_cell) was used to *Ptn* and *Ptprz1* brain cells expression under HFD condition (**Supplementary Figure 3**). The datasets are available from the corresponding author upon reasonable request.

## Author contributions

Conceptualization, funding acquisition, supervision, and project administration: MPR-A and GH. Investigation: HC-R, TF-B, MG-L, AZ, MGS-A and EG. Formal analysis and data curation: HC-R, MPR-A and GH. Data interpretation: HC-R, MPR-A, EG and GH. Writing - original draft: HC-R and GH. Writing - review & editing: HC-R, TF-B, MG-L, AZ, MGS-A, EG, MPR-A and GH. All authors have read and agreed to the published version of the manuscript.

## Acknowledgements

HC-R was supported by a fellowship from Fundación Universitaria San Pablo CEU-Santander. We thank our colleagues in the animal facility at the Universidad San Pablo CEU.

## Supplementary figure legends

**Supplementary figure 1. Representative experimental designs scheme.** Experimental design (A). Representative scheme of the novel object recognition (NOR) test (B).

**Supplementary Figure 2**: Food intake (g/mouse/day) of male and female *Ptn*^+/+^ STD, *Ptn*^+/+^ HFD, *Ptn*^−/−^ STD and *Ptn*^−/−^ HFD mice throughout the whole experiment.

**Supplementary Figure 3. High-fat diet feeding effects on *Ptn* and *Ptprz1* differential cell expression.** Single-cell analysis of *Ptn* and *Ptprz1* differential expression in mice samples grouped by clustering represented by t-distributed stochastic neighbour embedding (t-SNE) plot (A); Heatmap of *Ptn* and *Ptprz1* expression in different cell types from mice fed with a standard chow diet (STD) or with a high-fat diet (HFD) (n = 8657 cells) (B). \**p* < 0.05, \*\*\**p* < 0.001. Single-cell data was obtained from the Single Cell Portal published study “Multitissue Single-Cell Analysis Reveals Differential Tissue, Cellular, and Molecular Sensitivity Between Fructose and High Fat High Sucrose Diets”.

## REFERENCES

Ali, W., Ikram, M., Park, H. Y., Jo, M. G., Ullah, R., Ahmad, S., Abid, N. B., & Kim, M. O. (2020). Oral Administration of Alpha Linoleic Acid Rescues Aβ-Induced Glia-Mediated Neuroinflammation and Cognitive Dysfunction in C57BL/6N Mice. Cells, 9(3). 10.3390/cells9030667

Amet, L. E., Lauri, S. E., Hienola, A., Croll, S. D., Lu, Y., Levorse, J. M., Prabhakaran, B., Taira, T., Rauvala, H., & Vogt, T. F. (2001). Enhanced hippocampal long-term potentiation in mice lacking heparin-binding growth-associated molecule. Mol Cell Neurosci, 17(6), 1014–1024. 10.1006/mcne.2001.0998

Antunes, M., & Biala, G. (2012). The novel object recognition memory: neurobiology, test procedure, and its modifications. Cogn Process, 13(2), 93–110. 10.1007/s10339-011-0430-z

Arneson, D., Majid, S., Ahn, I. S., Kurt, Z., Diamante, G., Zhang, G., Gomez-Pinilla, F., & Yang, X. I. A. (2019). 1812-P: Multitissue Single-Cell Analysis Reveals Differential Tissue, Cellular, and Molecular Sensitivity between Fructose and High-Fat Diets. Diabetes, 68(Supplement_1), 1812-P. 10.2337/db19-1812-P

Cañeque-Rufo, H., Sánchez-Alonso, M. G., Zuccaro, A., Sevillano, J., Ramos-Álvarez, M. D. P., & Herradón, G. (2023). Pleiotrophin deficiency protects against high-fat diet-induced neuroinflammation: Implications for brain mitochondrial dysfunction and aberrant protein aggregation. Food Chem Toxicol, 172, 113578. 10.1016/j.fct.2022.113578

Castillo, X., Castro-Obregón, S., Gutiérrez-Becker, B., Gutiérrez-Ospina, G., Karalis, N., Khalil, A. A., Lopez-Noguerola, J. S., Rodríguez, L. L., Martínez-Martínez, E., Perez-Cruz, C., Pérez-Velázquez, J., Piña, A. L., Rubio, K., García, H. P. S., Syeda, T., Vanoye-Carlo, A., Villringer, A., Winek, K., & Zille, M. (2019). Re-thinking the Etiological Framework of Neurodegeneration. Front Neurosci, 13, 728. 10.3389/fnins.2019.00728

Cavaliere, G., Trinchese, G., Penna, E., Cimmino, F., Pirozzi, C., Lama, A., Annunziata, C., Catapano, A., Mattace Raso, G., Meli, R., Monda, M., Messina, G., Zammit, C., Crispino, M., & Mollica, M. P. (2019). High-Fat Diet Induces Neuroinflammation and Mitochondrial Impairment in Mice Cerebral Cortex and Synaptic Fraction. Front Cell Neurosci, 13, 509. 10.3389/fncel.2019.00509

Cope, E. C., & Gould, E. (2019). Adult Neurogenesis, Glia, and the Extracellular Matrix. Cell Stem Cell, 24(5), 690–705. 10.1016/j.stem.2019.03.023

Cornier, M. A., Dabelea, D., Hernandez, T. L., Lindstrom, R. C., Steig, A. J., Stob, N. R., Van Pelt, R. E., Wang, H., & Eckel, R. H. (2008). The metabolic syndrome. Endocr Rev, 29(7), 777–822. 10.1210/er.2008-0024

Crapser, J. D., Spangenberg, E. E., Barahona, R. A., Arreola, M. A., Hohsfield, L. A., & Green, K. N. (2020). Microglia facilitate loss of perineuronal nets in the Alzheimer’s disease brain. EBioMedicine, 58, 102919. 10.1016/j.ebiom.2020.102919

Dabke, K., Hendrick, G., & Devkota, S. (2019). The gut microbiome and metabolic syndrome. J Clin Invest, 129(10), 4050–4057. 10.1172/jci129194

de Bem, A. F., Krolow, R., Farias, H. R., de Rezende, V. L., Gelain, D. P., Moreira, J. C. F., Duarte, J., & de Oliveira, J. (2020). Animal Models of Metabolic Disorders in the Study of Neurodegenerative Diseases: An Overview. Front Neurosci, 14, 604150. 10.3389/fnins.2020.604150

Eill, G. J., Sinha, A., Morawski, M., Viapiano, M. S., & Matthews, R. T. (2020). The protein tyrosine phosphatase RPTPζ/phosphacan is critical for perineuronal net structure. J Biol Chem, 295(4), 955–968. 10.1074/jbc.RA119.010830

Fernández-Calle, R., Galán-Llario, M., Gramage, E., Zapatería, B., Vicente-Rodríguez, M., Zapico, J. M., de Pascual-Teresa, B., Ramos, A., Ramos-Álvarez, M. P., Uribarri, M., Ferrer-Alcón, M., & Herradón, G. (2020). Role of RPTPβ/ζ in neuroinflammation and microglia-neuron communication. Sci Rep, 10(1), 20259. 10.1038/s41598-020-76415-5

Fernández-Calle, R., Vicente-Rodríguez, M., Gramage, E., Pita, J., Pérez-García, C., Ferrer-Alcón, M., Uribarri, M., Ramos, M. P., & Herradón, G. (2017). Pleiotrophin regulates microglia-mediated neuroinflammation. J Neuroinflammation, 14(1), 46. 10.1186/s12974-017-0823-8

Galán-Llario, M., Gramage, E., García-Guerra, A., Torregrosa, A. B., Gasparyan, A., Navarro, D., Navarrete, F., García-Gutiérrez, M. S., Manzanares, J., & Herradón, G. (2024). Adolescent intermittent ethanol exposure decreases perineuronal nets in the hippocampus in a sex dependent manner: Modulation through pharmacological inhibition of RPTPβ/ζ. Neuropharmacology, 247, 109850. 10.1016/j.neuropharm.2024.109850

Galán-Llario, M., Rodríguez-Zapata, M., Gramage, E., Vicente-Rodríguez, M., Fontán-Baselga, T., Ovejero-Benito, M. C., Pérez-García, C., Carrasco, J., Moreno-Herradón, M., Sevillano, J., Ramos-Álvarez, M. P., Zapico, J. M., de Pascual-Teresa, B., Ramos, A., & Herradón, G. (2023). Receptor protein tyrosine phosphatase β/ζ regulates loss of neurogenesis in the mouse hippocampus following adolescent acute ethanol exposure. Neurotoxicology, 94, 98–107. 10.1016/j.neuro.2022.11.008

Gray, E., Thomas, T. L., Betmouni, S., Scolding, N., & Love, S. (2008). Elevated matrix metalloproteinase-9 and degradation of perineuronal nets in cerebrocortical multiple sclerosis plaques. J Neuropathol Exp Neurol, 67(9), 888–899. 10.1097/NEN.0b013e318183d003

Härtig, W., Derouiche, A., Welt, K., Brauer, K., Grosche, J., Mäder, M., Reichenbach, A., & Brückner, G. (1999). Cortical neurons immunoreactive for the potassium channel Kv3.1b subunit are predominantly surrounded by perineuronal nets presumed as a buffering system for cations. Brain Res, 842(1), 15–29. 10.1016/s0006-8993(99)01784-9

Herradón, G., & Ezquerra, L. (2009). Blocking receptor protein tyrosine phosphatase beta/zeta: a potential therapeutic strategy for Parkinson’s disease. Curr Med Chem, 16(25), 3322–3329. 10.2174/092986709788803240

Herradón, G., & Pérez-García, C. (2014). Targeting midkine and pleiotrophin signalling pathways in addiction and neurodegenerative disorders: recent progress and perspectives. Br J Pharmacol, 171(4), 837–848. 10.1111/bph.12312

Herradon, G., Ramos-Alvarez, M. P., & Gramage, E. (2019). Connecting Metainflammation and Neuroinflammation Through the PTN-MK-RPTPβ/ζ Axis: Relevance in Therapeutic Development. Front Pharmacol, 10, 377. 10.3389/fphar.2019.00377

Jakovljević, A., Tucić, M., Blažiková, M., Korenić, A., Missirlis, Y., Stamenković, V., & Andjus, P. (2021). Structural and Functional Modulation of Perineuronal Nets: In Search of Important Players with Highlight on Tenascins. Cells, 10(6). 10.3390/cells10061345

Liu, L., Li, T., Chang, J., Xia, X., & Ju, J. (2024). Microglia inversely regulate the level of perineuronal nets with the treatment of lipopolysaccharide and valproic acid. Neurosci Lett, 842, 137992. 10.1016/j.neulet.2024.137992

Maeda, N., Hamanaka, H., Shintani, T., Nishiwaki, T., & Noda, M. (1994). Multiple receptor-like protein tyrosine phosphatases in the form of chondroitin sulfate proteoglycan. FEBS Lett, 354(1), 67–70. 10.1016/0014-5793(94)01093-5

Migues, P. V., Liu, L., Archbold, G. E., Einarsson, E., Wong, J., Bonasia, K., Ko, S. H., Wang, Y. T., & Hardt, O. (2016). Blocking Synaptic Removal of GluA2-Containing AMPA Receptors Prevents the Natural Forgetting of Long-Term Memories. J Neurosci, 36(12), 3481–3494. 10.1523/jneurosci.3333-15.2016

Morawski, M., Brückner, M. K., Riederer, P., Brückner, G., & Arendt, T. (2004). Perineuronal nets potentially protect against oxidative stress. Exp Neurol, 188(2), 309–315. 10.1016/j.expneurol.2004.04.017

Ng, T. P., Feng, L., Nyunt, M. S., Feng, L., Gao, Q., Lim, M. L., Collinson, S. L., Chong, M. S., Lim, W. S., Lee, T. S., Yap, P., & Yap, K. B. (2016). Metabolic Syndrome and the Risk of Mild Cognitive Impairment and Progression to Dementia: Follow-up of the Singapore Longitudinal Ageing Study Cohort. JAMA Neurol, 73(4), 456–463. 10.1001/jamaneurol.2015.4899

Picone, P., Di Carlo, M., & Nuzzo, D. (2020). Obesity and Alzheimer’s disease: Molecular bases. Eur J Neurosci, 52(8), 3944–3950. 10.1111/ejn.14758

Pugazhenthi, S., Qin, L., & Reddy, P. H. (2017). Common neurodegenerative pathways in obesity, diabetes, and Alzheimer’s disease. Biochim Biophys Acta Mol Basis Dis, 1863(5), 1037–1045. 10.1016/j.bbadis.2016.04.017

Rebelos, E., Rinne, J. O., Nuutila, P., & Ekblad, L. L. (2021). Brain Glucose Metabolism in Health, Obesity, and Cognitive Decline-Does Insulin Have Anything to Do with It? A Narrative Review. J Clin Med, 10(7). 10.3390/jcm10071532

Reichelt, A. C., Hare, D. J., Bussey, T. J., & Saksida, L. M. (2019). Perineuronal Nets: Plasticity, Protection, and Therapeutic Potential. Trends Neurosci, 42(7), 458–470. 10.1016/j.tins.2019.04.003

Rodríguez-Matellán, A., Alcazar, N., Hernández, F., Serrano, M., & Ávila, J. (2020). In Vivo Reprogramming Ameliorates Aging Features in Dentate Gyrus Cells and Improves Memory in Mice. Stem Cell Reports, 15(5), 1056–1066. 10.1016/j.stemcr.2020.09.010

Rossier, J., Bernard, A., Cabungcal, J. H., Perrenoud, Q., Savoye, A., Gallopin, T., Hawrylycz, M., Cuénod, M., Do, K., Urban, A., & Lein, E. S. (2015). Cortical fast-spiking parvalbumin interneurons enwrapped in the perineuronal net express the metallopeptidases Adamts8, Adamts15 and Neprilysin. Mol Psychiatry, 20(2), 154–161. 10.1038/mp.2014.162

Santos, C. Y., Snyder, P. J., Wu, W. C., Zhang, M., Echeverria, A., & Alber, J. (2017). Pathophysiologic relationship between Alzheimer’s disease, cerebrovascular disease, and cardiovascular risk: A review and synthesis. Alzheimers Dement (Amst), 7, 69–87. 10.1016/j.dadm.2017.01.005

Sevillano, J., Sánchez-Alonso, M. G., Zapatería, B., Calderón, M., Alcalá, M., Limones, M., Pita, J., Gramage, E., Vicente-Rodríguez, M., Horrillo, D., Medina-Gómez, G., Obregón, M. J., Viana, M., Valladolid-Acebes, I., Herradón, G., & Ramos-Álvarez, M. P. (2019). Pleiotrophin deletion alters glucose homeostasis, energy metabolism and brown fat thermogenic function in mice. Diabetologia, 62(1), 123–135. 10.1007/s00125-018-4746-4

Sinha, A., Kawakami, J., Cole, K. S., Ladutska, A., Nguyen, M. Y., Zalmai, M. S., Holder, B. L., Broerman, V. M., Matthews, R. T., & Bouyain, S. (2023). Protein-protein interactions between tenascin-R and RPTPζ/phosphacan are critical to maintain the architecture of perineuronal nets. J Biol Chem, 299(8), 104952. 10.1016/j.jbc.2023.104952

Suttkus, A., Morawski, M., & Arendt, T. (2016). Protective Properties of Neural Extracellular Matrix. Mol Neurobiol, 53(1), 73–82. 10.1007/s12035-014-8990-4

Tewari, B. P., Chaunsali, L., Prim, C. E., & Sontheimer, H. (2022). A glial perspective on the extracellular matrix and perineuronal net remodeling in the central nervous system. Front Cell Neurosci, 16, 1022754. 10.3389/fncel.2022.1022754

Vicente-Rodríguez, M., Rojo Gonzalez, L., Gramage, E., Fernández-Calle, R., Chen, Y., Pérez-García, C., Ferrer-Alcón, M., Uribarri, M., Bailey, A., & Herradón, G. (2016). Pleiotrophin overexpression regulates amphetamine-induced reward and striatal dopaminergic denervation without changing the expression of dopamine D1 and D2 receptors: Implications for neuroinflammation. Eur Neuropsychopharmacol, 26(11), 1794–1805. 10.1016/j.euroneuro.2016.09.002

Wen, T. H., Afroz, S., Reinhard, S. M., Palacios, A. R., Tapia, K., Binder, D. K., Razak, K. A., & Ethell, I. M. (2018). Genetic Reduction of Matrix Metalloproteinase-9 Promotes Formation of Perineuronal Nets Around Parvalbumin-Expressing Interneurons and Normalizes Auditory Cortex Responses in Developing Fmr1 Knock-Out Mice. Cereb Cortex, 28(11), 3951–3964. 10.1093/cercor/bhx258

Więckowska-Gacek, A., Mietelska-Porowska, A., Wydrych, M., & Wojda, U. (2021). Western diet as a trigger of Alzheimer’s disease: From metabolic syndrome and systemic inflammation to neuroinflammation and neurodegeneration. Ageing Res Rev, 70, 101397. 10.1016/j.arr.2021.101397

Zuccaro, A., Zapatería, B., Sánchez-Alonso, M. G., Haro, M., Limones, M., Terrados, G., Izquierdo, A., Corrales, P., Medina-Gómez, G., Herradón, G., Sevillano, J., & Ramos-Álvarez, M. D. P. (2021). Pleiotrophin Deficiency Induces Browning of Periovarian Adipose Tissue and Protects against High-Fat Diet-Induced Hepatic Steatosis. Int J Mol Sci, 22(17). 10.3390/ijms22179261

