## Supplementary Figure 1 for "Pleiotrophin deletion prevents high-fat diet-induced cognitive impairment, glial responses, and alterations of the perineuronal nets in the hippocampus"

### Slide 1
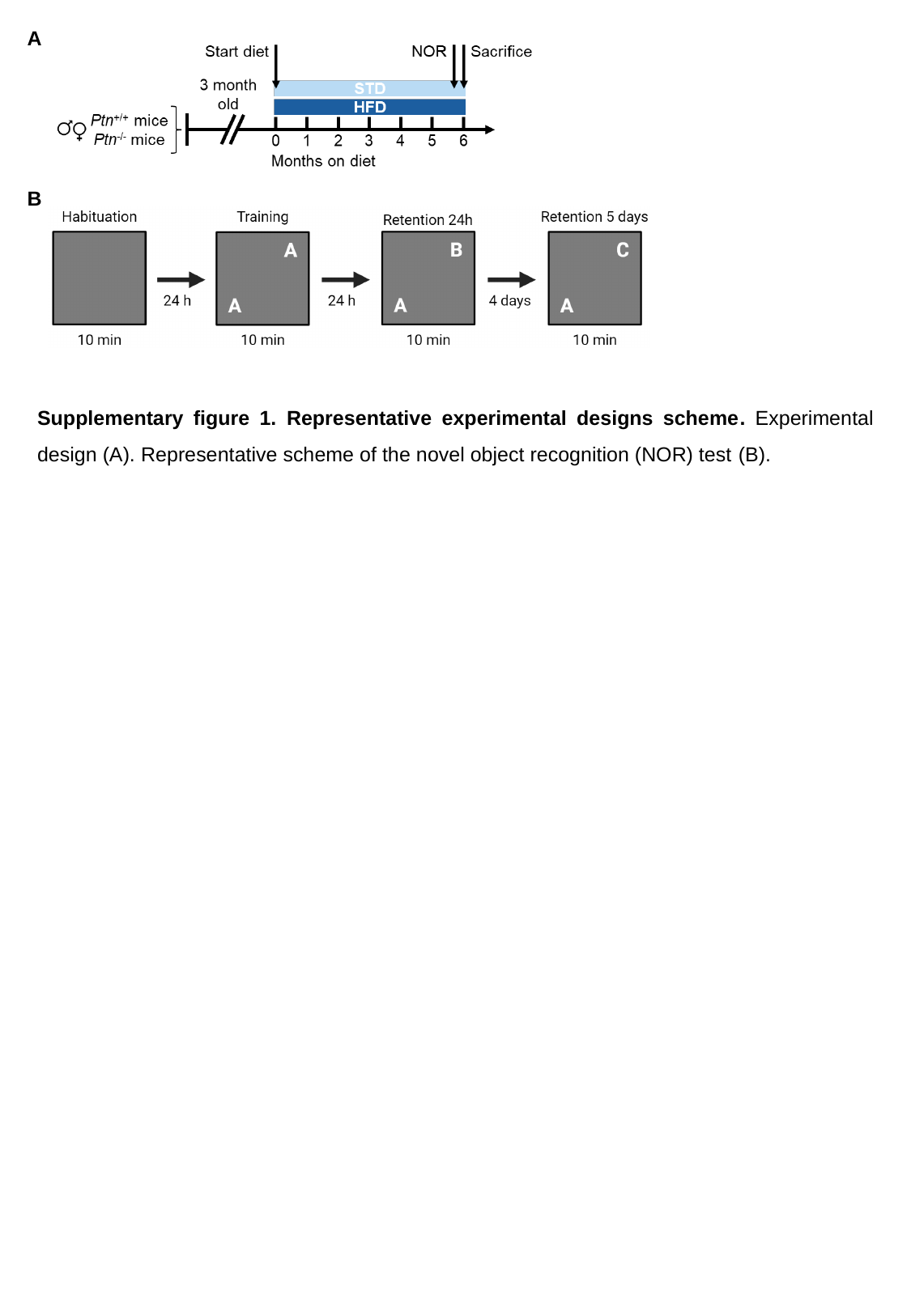

A
B
Supplementary figure 1. Representative experimental designs scheme. Experimental design (A). Representative scheme of the novel object recognition (NOR) test (B).
