## Supplementary Figure 2 for "Pleiotrophin deletion prevents high-fat diet-induced cognitive impairment, glial responses, and alterations of the perineuronal nets in the hippocampus"

### Slide 1
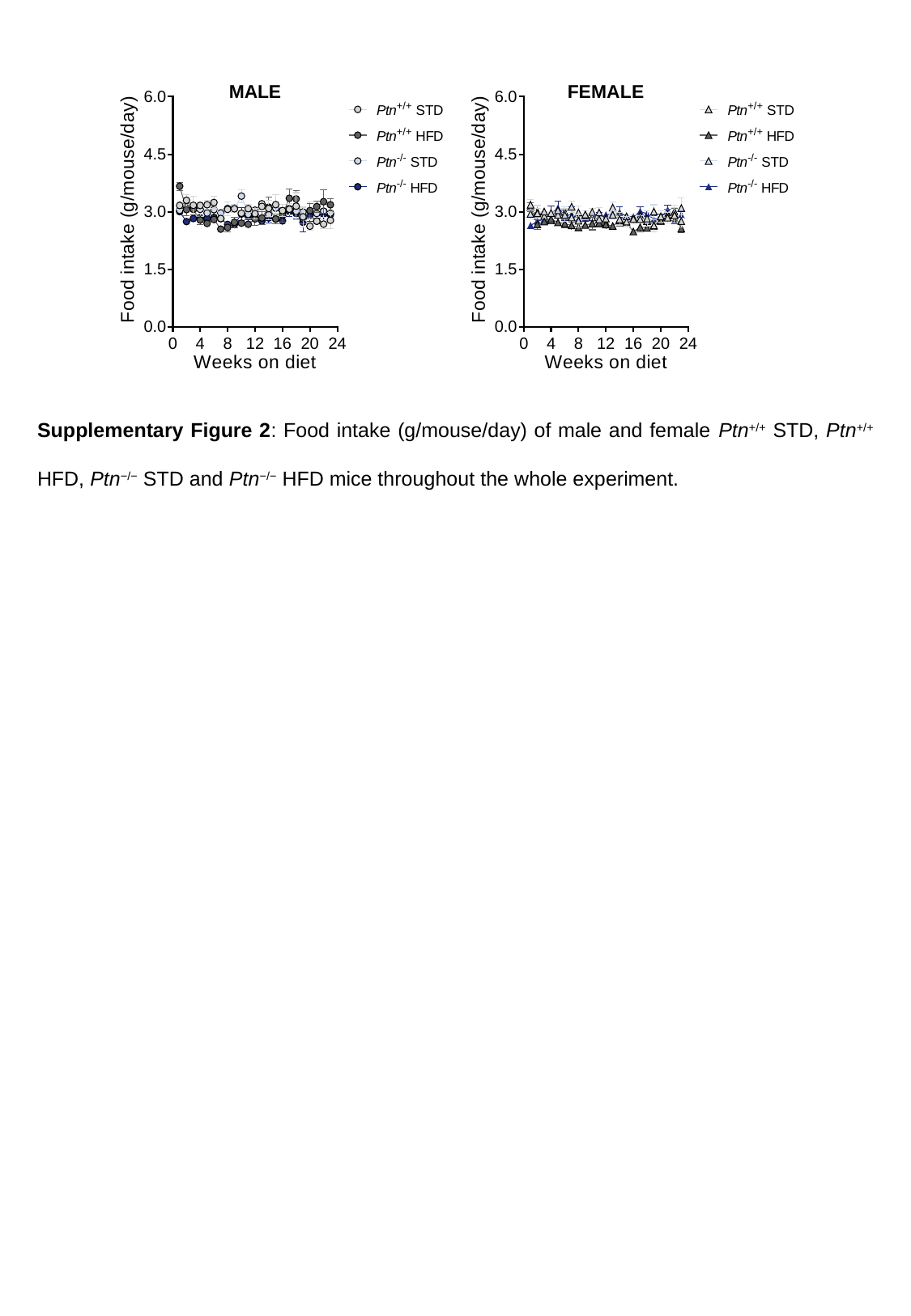

Supplementary Figure 2: Food intake (g/mouse/day) of male and female Ptn^+/+^ STD, Ptn^+/+^ HFD, Ptn^−/−^ STD and Ptn^−/−^ HFD mice throughout the whole experiment.
