## Supplementary Figure 3 for "Pleiotrophin deletion prevents high-fat diet-induced cognitive impairment, glial responses, and alterations of the perineuronal nets in the hippocampus"

### Slide 1
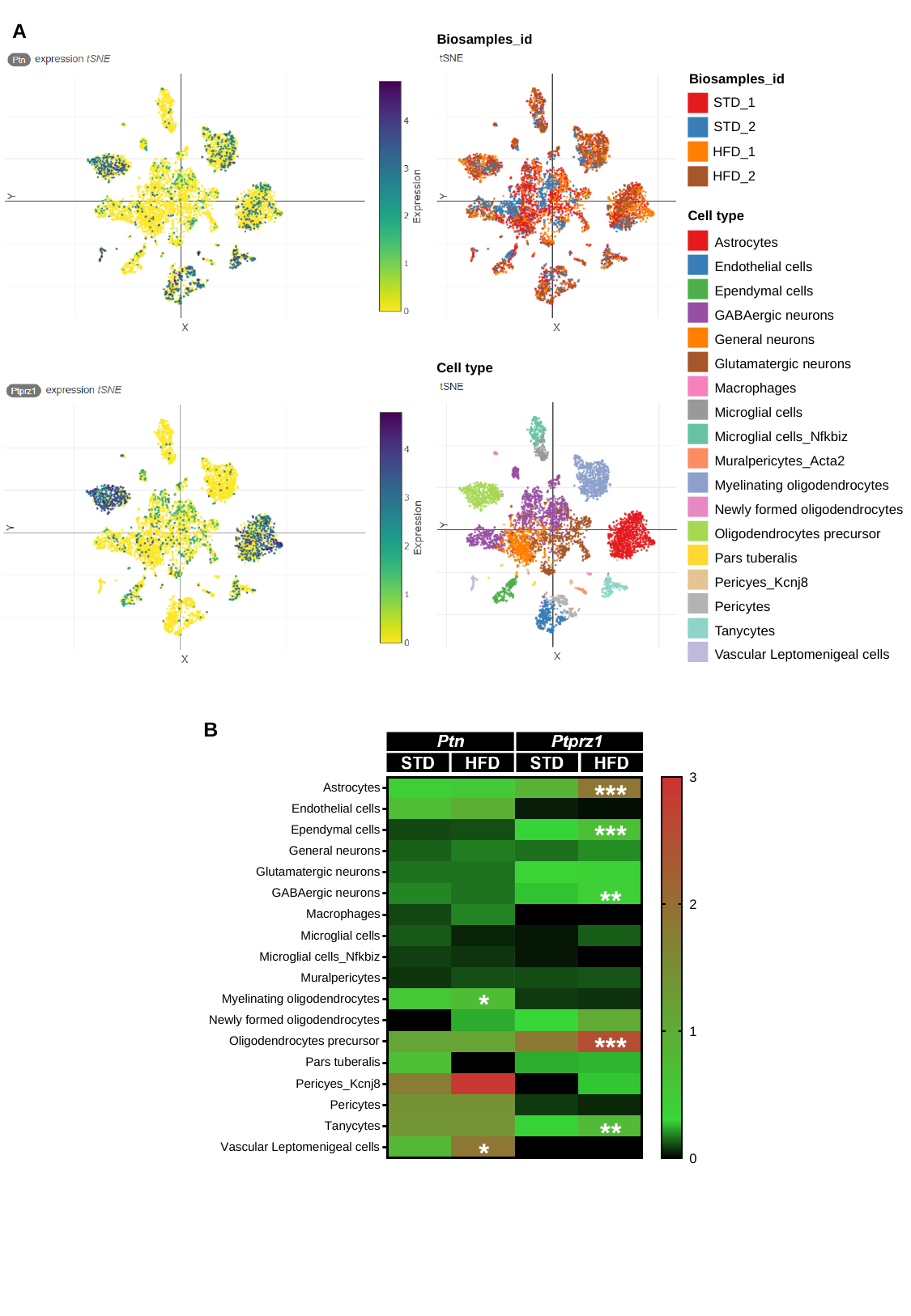

A
Biosamples_id
Biosamples_id
STD_1
STD_2
HFD_1
HFD_2
Cell type
Astrocytes
Endothelial cells
Ependymal cells
GABAergic neurons
General neurons
Glutamatergic neurons
Macrophages
Microglial cells
Microglial cells_Nfkbiz
Muralpericytes_Acta2
Myelinating oligodendrocytes
Newly formed oligodendrocytes
Oligodendrocytes precursor
Pars tuberalis
Pericyes_Kcnj8
Pericytes
Tanycytes
Vascular Leptomenigeal cells
Cell type
B

### Slide 2
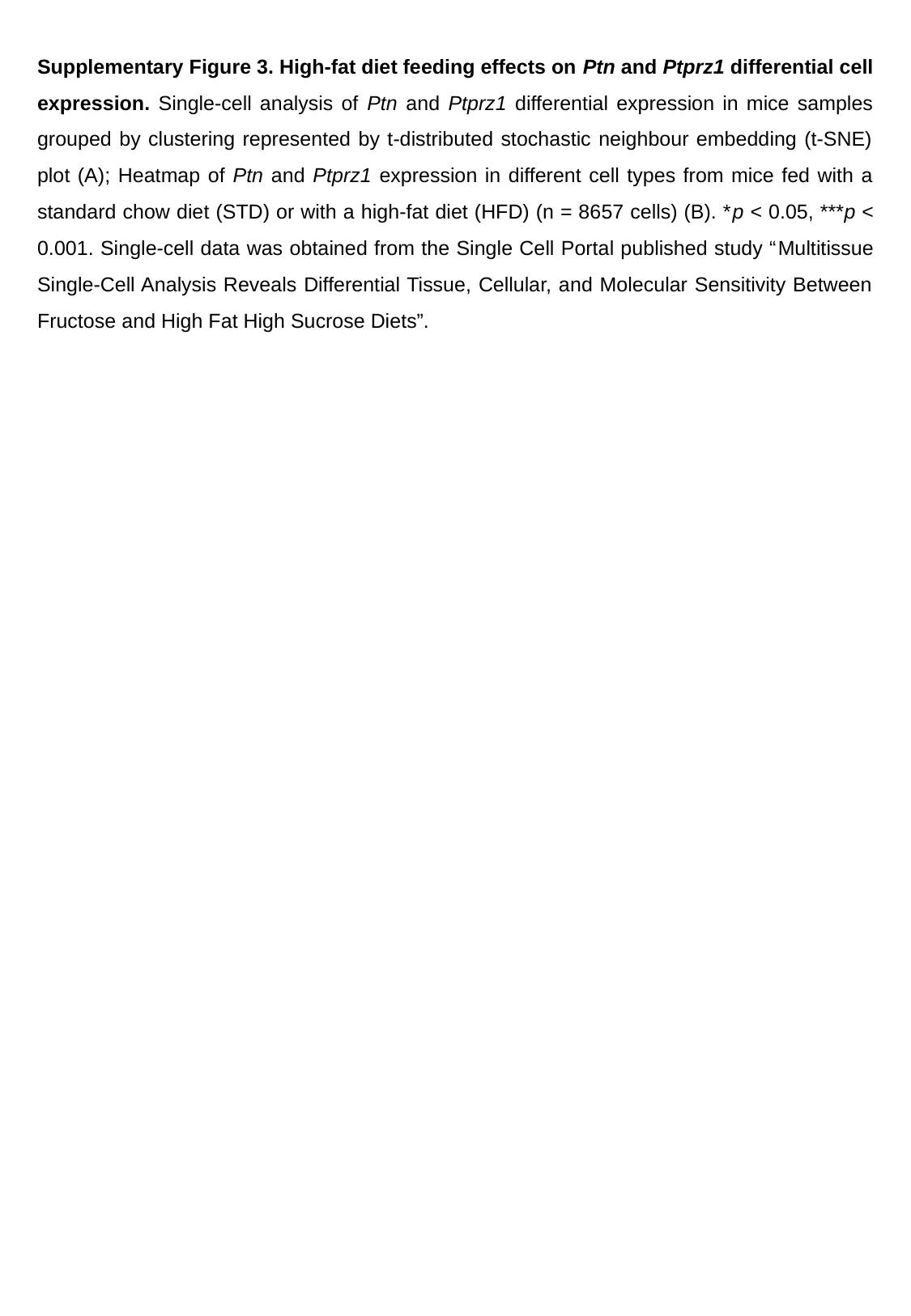

Supplementary Figure 3. High-fat diet feeding effects on Ptn and Ptprz1 differential cell expression. Single-cell analysis of Ptn and Ptprz1 differential expression in mice samples grouped by clustering represented by t-distributed stochastic neighbour embedding (t-SNE) plot (A); Heatmap of Ptn and Ptprz1 expression in different cell types from mice fed with a standard chow diet (STD) or with a high-fat diet (HFD) (n = 8657 cells) (B). *p < 0.05, ***p < 0.001. Single-cell data was obtained from the Single Cell Portal published study “Multitissue Single-Cell Analysis Reveals Differential Tissue, Cellular, and Molecular Sensitivity Between Fructose and High Fat High Sucrose Diets”.
